# SOX9 and SEMA7A regulate cell plasticity in the postpartum mammary gland with implications for breast cancer

**DOI:** 10.64898/2026.05.14.725155

**Authors:** Lauren M Cozzens, Brendan Hinckley, Alan M Elder, Veronica M Wessells, Sonali Jindal, Pepper J Schedin, Virginia F Borges, Traci R Lyons

## Abstract

Postpartum mammary gland involution is a coordinated process of cell death and remodeling that returns the tissue to a near pre-pregnant state following lactation and weaning. In models of postpartum breast cancer, defined as breast cancer diagnosed in women under age 45 and within 10 years of recent childbirth, involution induces durable phenotypes in breast tumor cells that promote progression and are associated with increased risk for therapeutic resistance, metastasis, and death in patients. SRY-Box Transcription Factor 9 (SOX9), a known regulator of mammary stem and progenitor cells, also promotes resistance to therapy and metastasis in breast cancers. Yet the contribution of SOX9 to the involution process is not well understood. We utilized single-cell RNA sequencing of mouse mammary glands during involution to delineate *Sox9*-expressing cell populations during lactation and involution. We found that *Sox9* mRNA is primarily expressed in luminal progenitor cells that are largely absent during lactation and present during early involution. We also reveal that *Sox9* is involved in a shift in cell state from lactational to non-lactational and is expressed in the surviving cells during involution. Prior work revealed that Semaphorin-7a (SEMA7A) also promotes cancer stem cell and pro-survival phenotypes in luminal progenitor cells during involution, and we observe a population of luminal progenitor cells that co-expresses *Sox9* and *Sema7a* during involution. Mechanistically, we demonstrate that knockdown of *Sox9* in cultured mammary epithelial cells results in increased SEMA7A expression, mesenchymal phenotypes, and loss of lactogenic differentiation capacity, identifying a potential regulatory axis where SOX9 balances SEMA7A expression in normal mammary epithelium and that disruption of this balance results in a dedifferentiated state that resembles mesenchymal cells. We validated a spatial relationship between SOX9 and SEMA7A proteins in a unique set of breast tissue samples from healthy human donors to show co-expression during early involution. In breast cancer datasets, we observe elevated expression of *SOX9* and *SEMA7A* in triple-negative breast cancers, as well as in the mesenchymal subtype of triple-negative breast cancers, suggesting disruption of this regulatory axis in breast cancer. Finally, we observe that co-expression increases metastatic risk in both estrogen receptor-negative and -positive breast cancers. Collectively, these findings define a novel SOX9–SEMA7A relationship in healthy mammary tissues and illustrate how studies of normal progenitor cell phenotypes can delineate cellular mechanisms that contribute to breast tumor progression.

## INTRODUCTION

Breast cancer diagnosed in women under age 45 and within 5-10 years of their most recent childbirth is defined as postpartum breast cancer (PPBC) based on the high rates of metastasis consistently observed in this patient population compared to age-matched nulliparous women with breast cancer^1–11^. Despite advances in understanding mechanisms that drive PPBCs, the underlying risk associated with recent pregnancy and the aggressive nature of PPBC remains incompletely understood. Preclinical studies identify cellular mechanisms associated with postpartum mammary gland involution – the tissue remodeling process that returns the gland to a pre-pregnant state – as drivers of PPBC progression^12–21^. Such studies have also led to the identification of potential biomarkers and novel therapeutic strategies for PPBC. Moreover, mechanisms uncovered in PPBC have provided insights into breast cancer progression more broadly and offer new perspectives on developmental pathways that merit further investigation.

The mammary gland undergoes the majority of its structural maturation postnatally and is organized as a branching ductal network terminating in lobuloalveolar units specialized for milk synthesis and secretion^22–25^. Within these structures, two mammary epithelial cell (MEC) lineages exist^26–28^, including luminal cells that line the ductal/alveolar lumen and basal cells that encase luminal cells and engage with the basement membrane^26,28,29^. During pregnancy, the luminal cells undergo expansion during alveologenesis, which is followed by differentiation to support milk production^30,31^. In unison, the basal/myoepithelial cells mature to acquire contractile phenotypes that enable milk ejection^31,32^. Mammary stem and progenitor cells enable epithelial regeneration throughout reproductive cycles and give rise to both the luminal and basal progenitor cells (LPCs and BPCs), which ultimately become the terminally differentiated luminal and basal cells^27,29,33–36^. While LPCs and BPCs in the adult mammary gland are largely considered lineage-restricted, emerging evidence indicates that both populations are multipotent in the adult gland^29,37,38^.

Postpartum mammary gland involution is initiated by milk stasis following weaning^39^. This involution process is characterized by coordinated cell death^39,40^, epithelial cell reprogramming^41^, and extracellular matrix remodeling^42^. Owing to limited access to healthy human breast tissue from the immediate post-weaning period, involution has been primarily defined through rodent models, which exhibit two well-characterized phases^43,44^. The first phase begins within hours of weaning and has a duration of two days^44^. During this phase, milk-producing luminal epithelial cells within the alveoli undergo cell death, while surrounding basal/myoepithelial cells remain intact to preserve gland structure^40,43–51^. The second phase occurs during a 3–6-day period when additional cell death, basement membrane breakdown, extracellular matrix remodeling, adipocyte repopulation, and re-organization of basal/myoepithelial cells occur^43–45,49,51,52^. Although alveolar collapse and tissue remodeling are necessary for involution, a subset of parity-associated MECs persists in the mammary gland after involution and expands with each pregnancy-lactation-involution cycle^53–56^. However, what factors govern the survival of epithelial cell populations to support subsequent lactation is not completely understood.

Mammary LPCs maintain stem-like characteristics, including self-renewal, plasticity^57^, and motility^58^, which render them susceptible to oncogenic transformation. In support of this, aberrant LPCs are proposed as cells of origin for basal-like breast cancers^29,57,59–61^. LPCs have been studied extensively across multiple stages of mammary gland development^33,62–68^; however, the mechanisms that regulate LPCs *during* involution are a continuing area of investigation. Given that involution promotes tumor progression, and evidence that LPCs persist following involution^33,38,63^, we hypothesize that aberrant LPCs may contribute to PPBC.

SRY-box transcription factor 9 (SOX9) and Semaphorin 7a (SEMA7A) promote progenitor-like phenotypes in MECs^66,69–71^. Specifically, studies have revealed that SOX9 is a transcriptional regulator of stem/progenitor cells in adult tissues^70,72^, that maintains stem-like and regenerative states in LPCs^69–73^ to enable their contribution to alveolar luminal lineages during pregnancy^74,75^. However, although it is recognized that SOX9^+^ cells persist after involution^75^, the expression dynamics and function of SOX9 *during* involution remain unclear.

Complementary studies have shown that SEMA7A, a GPI membrane-linked signaling protein that can be shed and signals through integrins^76,77^, is expressed on most of the live LPCs at the start of the second phase of involution^66^, and *Sema7a* mRNA expression was first identified in the mammary gland during the first phase of involution^45^. Consistent with a role in promoting maintenance of LPCs during involution, *Sema7a* knock-out (KO) mice exhibit decreased LPCs in the second phase of involution compared to wild-type (WT) mice^66^. Furthermore, studies of SEMA7A in mammary tissues during both puberty and involution identified roles for SEMA7A in non-epithelial cells^12,13,77,78^. *In vitro* mechanistic studies in cultured MECs revealed promotion of resistance to cell death by SEMA7A via alpha 6 and beta1 integrins (ITGA6+B1), including in conditions that resemble loss of attachment to the basement membrane, which occurs during the remodeling phase of involution^18,79^.

SOX9 promotes cancer stem cell (CSC) phenotypes, epithelial–mesenchymal transition/plasticity (EMT/P), and tumor progression in human cell lines and rodent models of breast cancer^69,70,74,80^. SOX9 expression is also elevated in basal-like tumors from patients^80^, which are thought to arise from aberrant LPC cells^59,60,81^. In rodent models of PPBC, which include postpartum and non-postpartum hosts, SEMA7A promotes tumor growth, EMT/P, CSC phenotypes, immune evasion, therapy resistance, and metastasis^12,13,66,77,78,82–85^. In PPBC patients, elevated SEMA7A protein is also an identifiable risk factor for recurrence^86^, and *SEMA7A* mRNA expression is associated with increased risk for progression across all breast cancer subtypes^82^.

Based on the body of evidence linking SOX9 and SEMA7A to progenitor cells and breast cancer, we hypothesized that SOX9, like SEMA7A, is expressed in MECs during postpartum involution and may be co-opted by PPBCs as well as all breast cancers that exhibit high expression. Using single-cell RNA-sequencing (scRNA-seq), we reveal that *Sox9* mRNA was expressed in luminal and basal MECs and specifically enriched in LPC populations during involution. We also reveal that SOX9 is expressed in non-apoptotic cells during early involution and persists after involution is complete. In contrast, *Sema7a* mRNA is expressed in a small subset of luminal populations during involution that are spatially situated between mature luminal and progenitor cell states. However, expression of mRNA for the SEMA7A receptors, *Itga6* and *Itgb1*, is abundant in LPCs and across basal and luminal MECs. In this small subset of *Sema7a*-expressing LPCs, we also observe dual expression of *Sox9*. Co-expression of SEMA7A and SOX9 protein in the early post-wean period was also validated for the first time in human tissues, using a cohort of breast tissues from healthy donors with known parity or involution status. To establish a relationship between SOX9 and SEMA7A in normal MECs, we utilized *in vitro* models of MEC lactogenic differentiation and cell death (LacDiff/Inv) to show that SEMA7A and SOX9 proteins both increase during the LacDiff/Inv phases. Mechanistically, we reveal that a consequence of *Sox9* knockdown (KD) in cultured MECs is increased expression of SEMA7A protein, gain of mesenchymal cell phenotypes, and loss of differentiation capacity, which was phenocopied by overexpression (OE) of SEMA7A. Finally, using publicly available breast cancer datasets, we reveal upregulation of *SOX9* and *SEMA7A* mRNAs in triple-negative breast cancers (TNBCs) compared to non-TNBC tumors and to normal breast tissue. Further, survival analysis demonstrates a significant decrease in distant metastasis-free survival (DMFS) in both estrogen receptor negative (ER-) and positive (ER+) breast cancers with co-high-expression of *SOX9* and *SEMA7A* that is not observed with *high SOX9* alone and is more significant than *high SEMA7A* alone. Collectively, our results suggest that SOX9 and SEMA7A are predominantly expressed on distinct cell populations in the mammary gland, but co-expression exists in a subset of cells in mouse and human tissues during the tumor promotional period of involution, and this co-expression may be co-opted by breast tumors to drive alterations in cellular plasticity and poor prognosis.

## RESULTS

### Single-cell sequencing and immunohistochemistry reveal *Sox9* expression in surviving murine mammary cells during postpartum involution

To characterize the expression of *Sox9* mRNA in MECs during involution, compared to lactation, we conducted scRNA-seq on cells isolated from mammary glands of age-matched mice post-parturition at lactation day 10 (Lac10) and at 3 days post force wean (Inv3) (**Supplemental Fig. 1A**). Lac10 represents peak lactation, occurring at ∼10 days post-parturition and prior to natural wean, and Inv3 represents the start of the second phase of involution. We fluorescence-activated cell sorted (FACS) the samples to isolate non-immune (EpCAM+CD45-) and immune cells (CD45+), for downstream analysis (**Supplemental Fig. 1A+B**). From the non-immune cells, we identified 8 unique clusters from Lac10 and Inv3 (**Fig. 1A**, **Supplemental Fig. 2A**). The MEC populations identified included luminal progenitor (LPCs), mature luminal, basal myoepithelial, and alveolar luminal. In the lactation sample, we observed a population of alveolar luminal cells that were reduced in involution samples (**Fig 1A**), and in the involution sample, the number of LPCs and luminal mature cells increased (**Fig. 1A**). We also observed basal myoepithelial populations from lactation that decreased in involution (**Fig. 1A**). *Sox9* was expressed in MECs from both lactation and involution samples, including LPCs, alveolar luminal, and basal myoepithelial cells at Lac10 (**Fig. 1A, Supplemental Fig. 2C**) and luminal mature, LPCs, and basal myoepithelial cells at Inv3 (**Fig. 1A; Supplemental Fig. 2C**). In addition, *Sox9* was among the top-enriched TFs in LPCs compared to all other populations (**Supplemental Fig. 2D**).

**Fig. 1.**
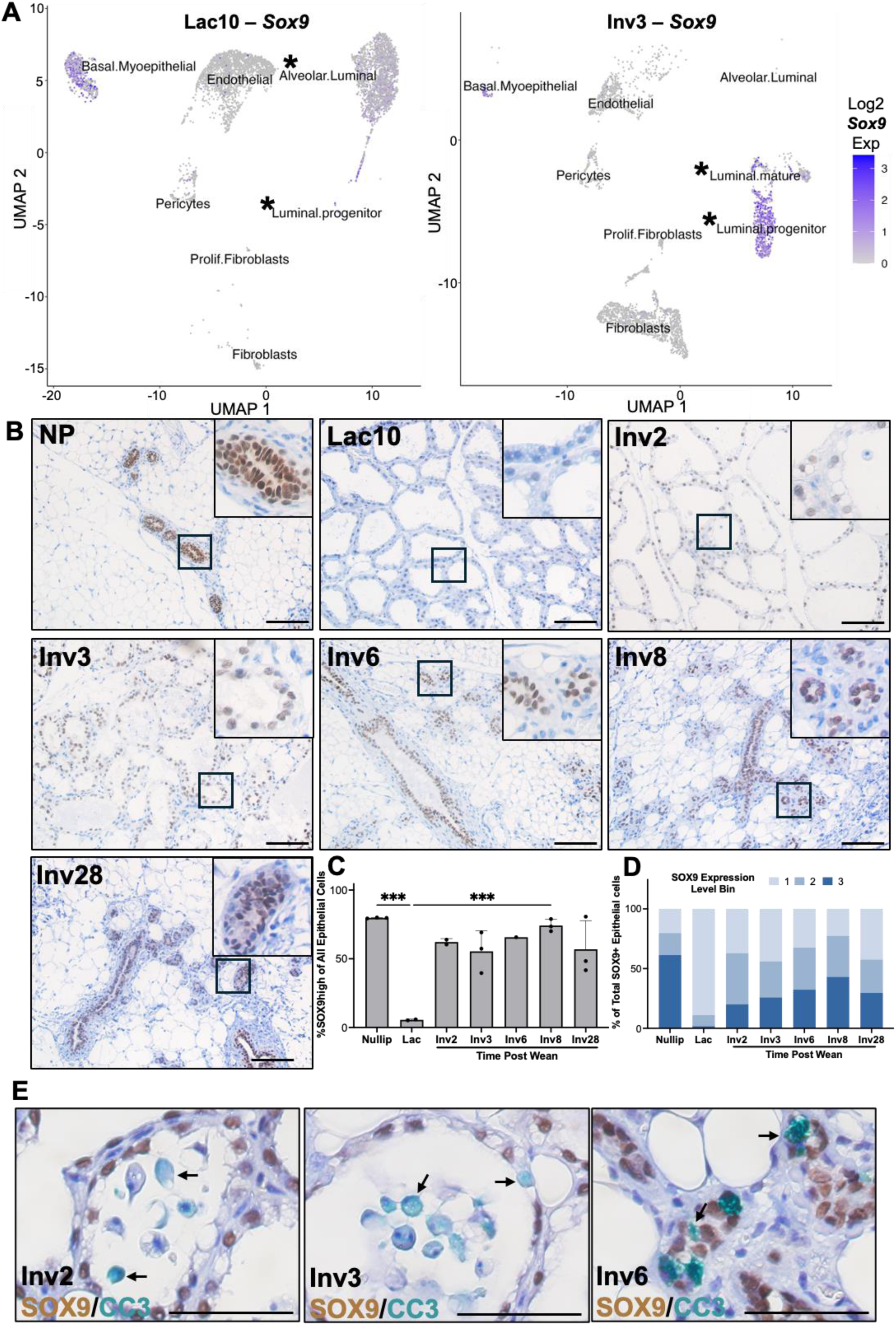
SOX9 is expressed in the murine mammary epithelium during involution in non-apoptotic cells. **A** UMAP visualization of non-immune populations from scRNA-seq of murine mammary glands from Lac day 10 (left) and Inv day 3 (right) with overlayed Log2 *Sox9* expression level. **B** Representative IHC images of murine mammary tissues isolated from WT C57BL/6J mice stained for SOX9 (brown) at the indicated reproductive timepoints. Scale bars: 1*00µm.* **B** Quantification of percent SOX9 high-expressing epithelial cells across Nulliparous (Nullip), Lactation (Lac), and Involution (Inv) time points. (Nullip n = 3, Lac10 n = 2, lnv2 n = 2, lnv3 n = 3, lnv6 n = 1, lnv8 n = 3, lnv28 n = 3). C Quantification of SOX9 expression level bins in epithelial cells across Nullip, Lac, and Inv time points (1 = low, 2 = medium, 3 = high). D Representative IHC images of mouse mammary tissues stained for SOX9 (brown) and CC3 (green). (lnv2 n = 2, lnv3 n = 3, lnv6 n = 1). Error bars represent the mean ± SD. Ordinary one-way ANOVA Sidak’s Multiple comparisons test, no asterisk p > 0.05, *\*\*\*p* ≤ 0.001.

To explore the role of SOX9 protein during the transition from lactation to involution and at additional involution timepoints, we conducted immunohistochemistry (IHC) staining for SOX9 protein in murine mammary tissues during involution, and nulliparous, lactation, and fully regressed (Inv28) samples (**Fig. 1B**). We found SOX9-expressing mammary epithelial cell nuclei from nulliparous animals, which decreased during lactation (**Fig. 1B-C**). Expression increased during involution, where we observed SOX9-positive (SOX9+) nuclei at involution days 2, 3, 6, and 8, which remained high at involution day 28, which is considered fully regressed (**Fig. 1B-C**). We also examined SOX9 staining intensity to show that the expression level also decreased during lactation, compared to nulliparous tissue, and returned in involution samples (**Fig. 1D**). To investigate the relationship between SOX9 and live versus apoptotic cells, dual-stained mammary tissues were examined for SOX9, when apoptosis of the epithelium is high (Inv days 2, 3, and 6), and cleaved-caspase-3 (CC3) as a marker of apoptosis. At Inv2 and 3 we observed distinct SOX9 expression in the epithelium in the alveoli lumens, with some CC3 expression in alveoli associated cells that were SOX9-negative (SOX9-), and CC3 expression in the apoptotic bodies shed into the lumen (**Fig. 1E, Supplemental Fig. 3A**). At Inv6, we observed cells that were both CC3 positive and negative in the alveoli but could not discern whether those cells were dual positive for SOX9. In addition, we observed very little Ki67 expression during involution, suggesting that the MECs present are mostly quiescent at these time points (**Supplemental Fig. 3A+B**). Given the high percentage of SOX9+ cells in alveolar structures and ducts post-involution, we suggest that SOX9+ cells do not undergo apoptosis during involution and therefore persist in the post-involution state. This result is similar to our previous studies of postpartum involution in mice, where SEMA7A surface protein expression was present on nearly all the live, non-apoptotic, LPCs during involution^66^.

### SOX9 and SEMA7A in LPCs and models of MEC differentiation and cell death reveal mechanisms of cellular reprogramming

Given the apparent relationship between SOX9 and SEMA7A in promoting LPC survival during involution, we examined *Sema7a* in our scRNASeq. Small subsets of *Sema7a*-expressing luminal and basal MECs were identified in the lactation and involution samples (**Fig. 2A; Supplemental Fig. 4A+B**). We also examined expression of known receptors for SEMA7A, including *Plexinc1*^87^ and *Itga6*+*b1*^66^, to show that *Plexinc1* expression was low, while *Itga6+b1* was co-expressed in all MEC subtypes (**Supplemental Fig. 4C+D**). Further analysis of the LPC populations during involution revealed that a small subset express *Sema7a* (yellow) and are spatially related to a subset of *Sox9-*expressing LPCs (blue) (**Fig. 1A**). These cells are positioned at the interface between luminal mature and progenitor cell states, and some cells within this cluster exhibited co-expression of *Sox9* and *Sema7a* (green) (**Fig. 1A**). Additional sub-clustering analysis of the LPC populations from both lactation and involution identified 4 unique clusters, which were predominantly derived from the Inv3 sample; co-expression of *Sema7a* and *Sox9* was restricted to LPC cluster 3 (**Supplemental Fig. 4E-G**). Gene Ontology Biological Process (GOBP) and Hallmarks of Cancer (Hallmark) pathway analysis of cluster 3 LPCs revealed enrichment for GOBP pathways relating to epithelial development, as well as ones related to non-epithelial cells, including angiogenesis and migration. In addition, hallmarks of cancer pathways relating to TNFa/NFkb, apoptosis, hypoxia, p53, TGF-*β* signaling, and cholesterol homeostasis were observed in cluster 3 (**Fig. 2B**).

**Fig. 2.**
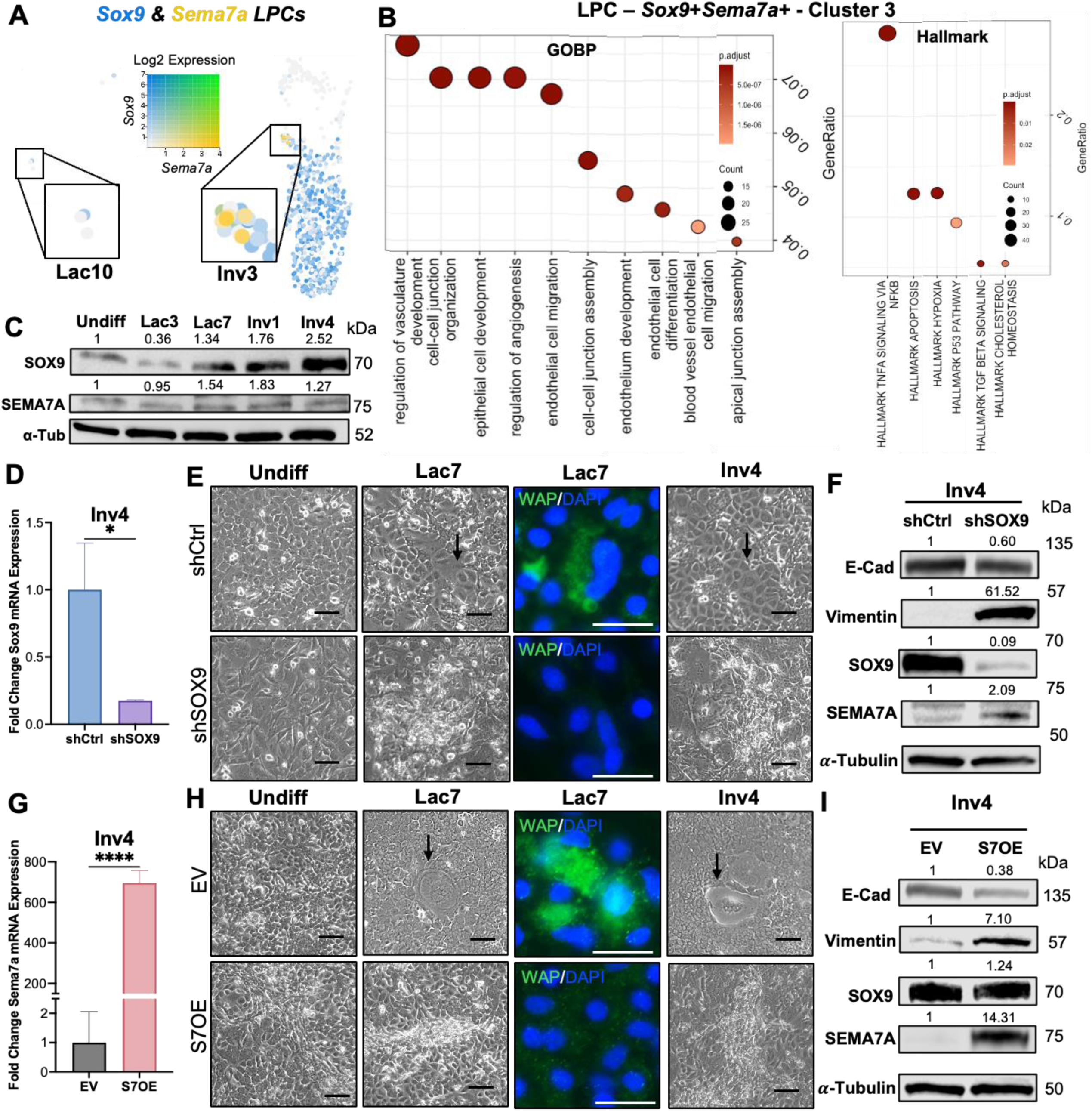
*Sox9* and *Sema7a* are co-expressed in a subset of LPCs during involution *in vivo* and affect MEC differentiation *in vitro.* **A** Close-up of LPC populations from scRNA-seq of Lac? (left) and lnv3 (right) samples overlayed with Log2 *Sox9 and Sema7a* co-expression levels. **B** Dot plots of significantly upregulated gene ontology biological process pathways (GOBP, left) and Hallmarks of Cancer pathway (Hallmarks, right) analysis for the LPC subcluster, where *Sox9* and *Sema7a* are both expressed (Cluster 3). **C** lmmunoblot showing protein levels of SOX9 and SEMA7A in undifferentiated (Undiff), early (Lac3), and late lactogenic differentiation (Lac7), and early (Inv1) and late involution (Inv4) in Wildtype (WT) HC-11 cell cultures (representative blot of n = 3 independent experiments). **D+G** qPCR for *Sox9* mRNA expression in HC-11 cells harboring an shCtrl or shSOX9 shRNA construct **(D),** or *Sema7a* mRNA expression in HC-11 cells harboring an EV or S7OE vector **(G),** represented as fold change mRNA expression relative to shCtrl or EV controls, respectively (representative plots from n = 2 independent experiments with technical triplicates). RNA was extracted on Inv day 4. E+H Brightfield images of HC-11 shCtrl+shSOX9 (E) and EV+S7OE **(H)** cultures across LacDiff/lnv, black arrows indicate dome formation and lmmunofluorescence of HC-11 shCtrl+shSOX9 and EV+S7OE cells for Whey Acetic Protein (WAP, green) on Lac day 7; nuclei stained with DAPI (blue). Scale bars = *20µm* (representative blot from n = 3 independent experiments). F+I lmmunoblot for SOX9, SEMA7A, E-Cadherin (E-Cad), and Vimentin protein levels in HC-11 shCtrl+shSOX9 (F) and EV+S7OE **(1)** cells. Alpha tubulin (a-Tubulin) was used as a loading control (representative blot from n = 3 independent experiments). All samples collected on lnv4. Error bars represent the mean ± SD. Quantification of immunoblots represented as fold change relative to control or undifferentiated cell cultures. Unpaired two-tailed t-test, *\*p* ≤ 0.05, *\*\*\*\*p* ≤ 0.0001.

To examine MEC phenotypes driven by SOX9 during lactation and involution, we utilized MEC *in vitro* models of LacDiff/Inv. Both HC-11 and EpH4 murine MECs were grown to confluence and treated with lactogenic hormones to promote differentiation (LacDiff), followed by hormone withdrawal to simulate involution (Inv) (**Supplemental Fig. 5A**). In these models, exposure to lactogenic hormones results in characteristics reminiscent of differentiated mammary MECs *in vivo,* evidenced by “dome” structures, which have been proposed as rudimentary gland-like structures^88^ that express milk proteins, including whey acidic protein (WAP) (**Supplemental Fig. 5B; 6A**). Dome formation and milk protein production were not observed in control cultures grown under the same conditions in the absence of lactogenic hormones (**Supplemental Fig. 7**). Partial collapse of the domes occurs upon hormone withdrawal, which is accompanied by increased cleaved caspase 3 activity, resembling involution (**Supplemental Fig. 5B+C; 6A+B**). We observed that both SOX9 and SEMA7A are endogenously expressed and are expressed at varying levels during both the lactation and involution phases. Specifically, both increase at the late LacDiff timepoint, and are further elevated at Inv day 1 (Inv1) (**Fig. 2C**). While SEMA7A expression appears transient, as evidenced by decreased levels at Inv4, SOX9 further increases at Inv4 (**Fig. 2C**), consistent with our previous studies in which SEMA7A was transiently expressed on MECs during involution^66^, and our current IHC analysis where SOX9 MEC expression remained high in late involution and fully regressed samples.

To understand the necessity of SOX9 in maintaining epithelial phenotypes throughout the LacDiff/Inv cycle, we engineered cells with stable shRNA-mediated KD of *Sox9* (shSOX9). We confirmed *Sox9* KD at the mRNA and protein levels, comparing KD cells to their non-targeting shRNA (shCtrl) controls (**Fig. 2D+F: Supplemental Fig. 5D+E, 6D+F, 8A+B**). We observed decreased dome formation alongside altered morphology and decreased WAP production, reminiscent of impaired LacDiff, in shSOX9 and cells compared to controls (**Fig. 2E; Supplemental Fig. 6E**). As a consequence of *Sox9* KD, upregulation of SEMA7A protein expression occurred at Inv4, which was confirmed at the mRNA level, and at the protein level in undifferentiated and Lac7 samples (**Fig. 2F**; **Supplemental Fig. 5E+H**, **6F**, **8B**).To model SEMA7A upregulation and examine how sustained SEMA7A expression impacts epithelial phenotypes, we engineered cells with stable overexpression of *Sema7a* (S7OE). We confirmed S7OE at the mRNA and protein levels, comparing all cells to their empty vector (EV) controls (**Fig. 2G+I: Supplemental Fig. 5F+G, 6D+F, 8C+D**). S7OE phenocopied Sox9 KD as evidenced by decreased dome formation and WAP production and altered epithelial morphology (**Fig. 2H; Supplemental Fig. 6E**). Thus, we evaluated the expression of E-Cadherin and Vimentin as representative markers of epithelial and mesenchymal phenotypes^89^, respectively, at Inv4. In both the *Sox9* KD and the S7OE cells, we observed downregulation of E-Cadherin expression and upregulation of Vimentin expression (**Fig. 2F+I**). Similar results were observed in undifferentiated (Undiff) cultures and in cultures harvested during LacDiff (**Supplemental Fig. 5E+G**). E-Cadherin downregulation and Vimentin upregulation were confirmed in EpH4 cultures, as well as with S7OE – albeit to a lesser extent (**Supplemental Fig. 6F, 8B+D**). In all cell states, S7OE did not affect SOX9 expression (**Fig. 2I, Supplemental Fig. 5G+H, 6F, 8D**).

### Disruption of SOX9 and overexpression of SEMA7A reveal hybrid epithelial-mesenchymal phenotypes via distinct molecular pathways

To quantify the morphologic changes observed with *Sox9* KD or S7OE, LacDiff/Inv cultures were stained with phalloidin to visualize the cytoskeleton and calculate cell aspect ratios (cell width/cell length), where an aspect ratio near 1 indicates a more cuboidal cell shape characteristic of epithelial cells, while an aspect ratio of < 1 corresponds to a more elongated mesenchymal-like morphology^90,91^. Our results indicate that both *Sox9* KD and S7OE resulted in aspect ratios significantly lower than their respective controls in both the Lac and Inv phases (**Fig. 3A-D, Supplemental Fig. 9A-D**), indicating a more mesenchymal morphology with *Sox9* KD and S7OE.

**Fig. 3.**
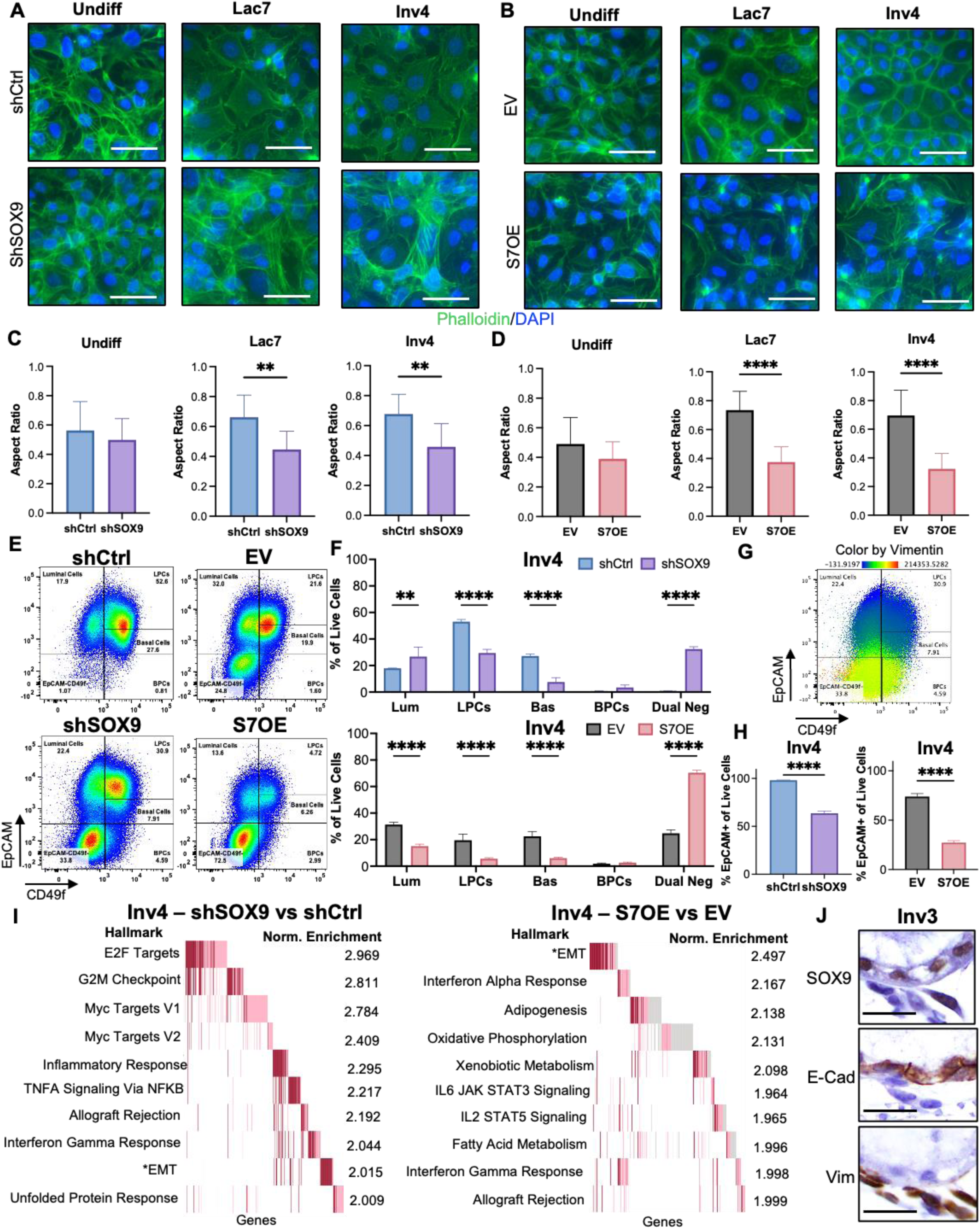
Loss of SOX9 or gain of SEMA7A promotes the emergence of a mesenchymal cell state in HC-11 models of involution. **A+B** Phalloidin stain images of HC-11 shCtrl+shSOX9 (A) and EV+S7OE **(B)** cultures across LacDiff/lnv (representative photos from n = 2 independent experiments). Scale bars = 20 *µ* m. C+D Aspect ratio calculations (width/length) from phalloidin staining in A+B (n = 10 cells/group). E Representative flow cytometry plots of EpCAM (y-axis) and CD49f (x-axis) staining in HC-11 cultures to visualize MEC subtypes, shCtrl+shSOX9 (left) and EV+S7OE (right). Numbers denote the percentage of that subtype of all live cells (representative plots of n = 3 independent experiments with technical triplicates). F Quantification of percent individual MEC subtypes from flow cytometry of all live cells in HC-11 shCtrl+shSOX9 cells (top) and EV+S7OE cells (bottom). G Example flow cytometry plot of EpCAM (y-axis) and CD49f (x-axis) staining, colored by Vimentin fluorescence intensity. H Quantification of percent EpCAM+ cells from flow cytometry of all live cells in HC-11 shCtrl+shSOX9 cells (left) and EV+S7OE cells (right). I Top 10 most significantly (adj. p-value) enriched Hallmark of Cancer (Hallmark) pathways from RNA-seq of HC-11 shSOX9 (left) or S7OE (right) lnv4 cultures compared to their associated controls, shCtrl and EV, respectively. Normalized enrichment values shown. *denotes epithelial to mesenchymal transition (EMT) as a top pathway for both cell lines. (n = 3 biological triplicates). J Representative images of serial sections of murine mammary glands from Inv day 2, 3, or 6 stained for SOX9, E-Cadherin, or Vimentin. Scale bars = *20µm.* Error bars represent the mean ± SD. C+H Unpaired two-tailed t-test. **F** Two-way ANOVA Sidak’s Multiple comparisons test. No asterisk p > 0.05, *\*\*p* ≤ 0.01, *\*\*\*\*p* ≤ 0.0001.

Utilizing a gating strategy like that used to distinguish mouse MEC subpopulations *in vivo*^57,81,92^, we performed flow cytometry to validate and extend our analysis of the shSOX9 and S7OE cells. We utilized EpCAM and CD49f to resolve static cell states in luminal cells (Lum - EpCAM^+^CD49f^-^), LPCs (EpCAM^+^CD49f^+^), basal cells (Bas - EpCAM^low/mid^CD49f^+^), BPCs (EpCAM^-^CD49f^+^), and dual negative cells (Dual Neg – EpCAM^-^CD49f^-^) isolated from Undiff, Lac7, and Inv4 cultures (**Supplementary Fig. 10**). Compared to controls, *Sox9* KD resulted in decreased LPCs and basal cells, which was also observed in the S7OE cells (**Fig. 3E+F, Supplemental Fig. 11A, B, E+F**). We also consistently observed that the EpCAM-CD49f-(Dual Neg), presumably a more mesenchymal population, increased upon *Sox9* KD or S7OE for both cell lines (**Fig. 3E+F, Supplementary Fig. 9E+F, 11A, B, E+F, 12A, B, E+F**). This dual negative population is typically excluded from primary MEC analysis of *ex vivo* mammary glands for which this gating strategy was developed, and is therefore understudied in the literature, where it is classified as “stromal cells” ^93,94^; this population was excluded from our scRNA-seq analysis presented in **Fig.1+2,** which partially explains why SEMA7A expression was rare. Importantly, as these are established MEC cultures that should not contain any true stromal cell populations, this dual negative population could represent an epithelial cell-derived mesenchymal-like cell state that is driven by SEMA7A. In support of this, we observe enrichment of the mesenchymal marker, vimentin, in the dual negative population (**Fig. 3G, Supplemental Fig. 9G, 11C+G, 12C+G**), which coincided with a decrease in the proportion of EpCAM+ (epithelial) cells (**Fig. 3H, Supplemental Fig. 9H, 11D+H, 12D+H**). We confirmed epithelial-mesenchymal transition/plasticity (EMT/EMP) by showing that *Snail* and *Zeb1* mRNAs, transcriptional regulators of mesenchymal cell programs^95^, are expressed at Inv4, with increased expression in *Sox9* KD or S7OE cells compared to controls (**Supplemental Fig. 13A**). To further analyze mesenchymal transcriptional programs, we performed bulk RNA-seq on HC-11 cultures at Inv4 and confirmed that mRNA expression of *Cdh1* (E-Cadherin) was decreased, while *Vim*, *Snai1* (Snail), and *Zeb1* were increased in shSOX9 or S7OE Inv4 cells compared to controls (**Supplemental Fig. 13B**). Expression of the EMT transcription factor *Snai2* (Slug) was decreased in shSOX9 KD cells but increased in S7OE cells compared to controls (**Supplemental Fig. 13B**). In addition, Hallmarks of Cancer pathway analysis revealed similar enrichment for EMT associated genes in both shSOX9 and S7OE Inv4 cultures relative to controls, as well as distinct gene set enrichments in each cell type, including cell cycle and Myc targets in shSOX9 cells and interferon response and metabolism in S7OE cells (**Fig. 3I**). This enrichment was also confirmed in EpH4 *Sox9* KD cultures compared to shCtrl cells (**Supplemental Fig. 9I**). To examine the functional impact of *Sox9* KD and S7OE at Inv4 we evaluated cell motility as a key phenotype of mesenchymal cells and observed that *Sox9* KD cells exhibited increased motility (**Supplemental Fig. 13C**). While this effect was not observed in S7OE cells compared to the controls it is noted the control cells in this assay were already more motile than shCtrl cells. We also stained serial sections of normal mammary tissue from mice during involution to show that SOX9 is primarily restricted to E-Cadherin-expressing epithelial cells, and that there is a close association of E-cadherin and vimentin-positive cells (**Fig. 3J; Supplemental Fig. 13D**).

### SOX9 and SEMA7A expression is conserved in human mammary tissues

To determine whether SOX9 and SEMA7A expression patterns are conserved in human breast tissue, we performed multi-spectral IHC (mIHC) on breast tissue donated by healthy women aged 20-45 years, where their parity status and timing of lactation/weaning were known^96^. This unique cohort included tissue from a University of Colorado Cancer Center study of weaning-induced breast involution, where women provided lactation history, establishing a set of post-wean samples between 0.5 and 6 months following cessation of lactation. Additional breast tissues from nulliparous and actively lactating women were obtained from the Komen Tissue Bank for comparison. Tissues were segmented into epithelial regions, which increased in lactation samples compared to nulliparous, and decreased in the post-weaning period (**Supplemental Fig. 14A+B**). We observed that the percentage of SOX9+ cells was high in the breast epithelium from nulliparous samples, decreased in lactation samples, and was re-established by 1 month post-wean (**Fig. 4A+B**). In addition, SOX9 mid-expressing cells remained higher in tissues with parity, compared to nulliparous, as late as 6 months post-wean (**Fig. 4C**). To evaluate SOX9+ epithelial populations further, we examined expression of cytokeratin 5 (CK5) to identify basal/myoepithelial cells, and cytokeratin 8 and 18 (CK8/18) to identify luminal cells^97^ and used co-expression to identify basal/luminal hybrid cells, which are thought to represent an intermediate subset of highly plastic cells^7,38,98–101^ also observed in breast cancers^102–104^ (**Fig. 4D, Supplemental Fig. 14C**). The overall proportion of hybrid cells did not change throughout the timepoints observed (**Supplemental Fig. 14D**). However, approximately 30-50% of SOX9+ cells were classified as luminal, about 10-40% were classified as basal, while about 30-40%were classified as basal/luminal hybrid cells and this pattern was maintained across samples (**Fig. 4D+E**). Alternatively, SOX9 expression was observed in large percentages of basal/luminal hybrid cells in both the nulliparous and post-wean epithelium (**Fig. 4F**).

**Fig. 4.**
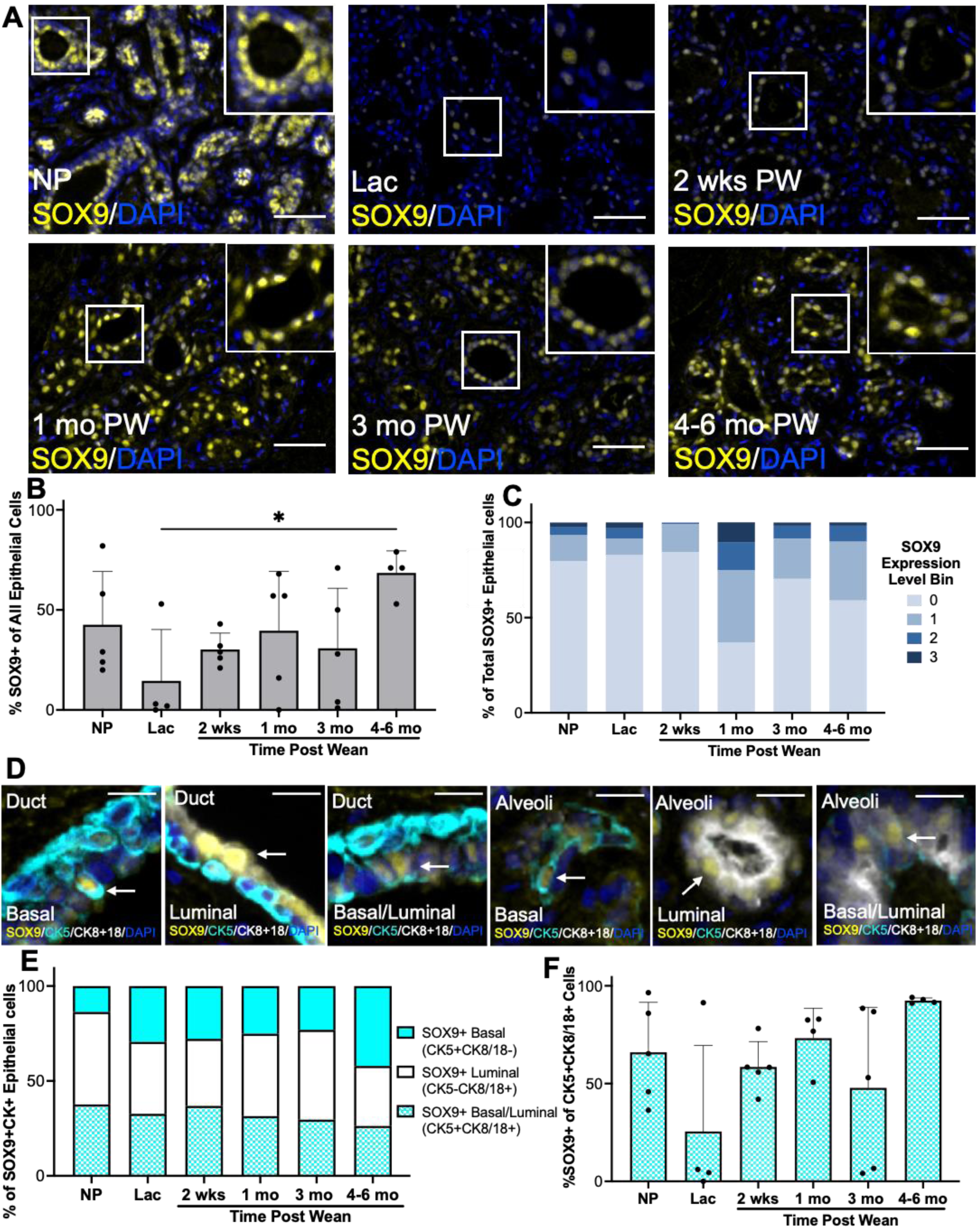
SOX9 expression in the human mammary epithelium is reduced duriing lactation and restored post-weaniing,, and marks cell populations exhibiting both basal and luminal features.. **A** Representative m1ulti-spectral IHC (mlHC) image of human mammary tissue stained for SOX9 (yellow) at indicated Nulliparous (Nullip/NP), Lactation (Lac), and Post-Wean (PW) time points; nuclei stained with DAPI (blue). Scale bars: 50 *µ* m. B Quantification of percent SOX9+ epithelial cells of all epithelial cells per tissue section. C Quantification and binning of percent SOX9 expression level from low (0) to high (5) of total SOX9+ epithelial cells per tissue section. **D** Representative mlHC imag1es of human mammary tissue stained for SOX9 (yellow), Cytokeratin5 (CK5 cyan), and Cytokeratin 8/18 (CK8+18 white), showing SOX9+ CK5+CK8/18-, Basal cells, CK5-CK8/18+ Luminal cells, and CK5+CK8/18+ Basal/Luminal hybrid cells; nuclei stained with DAPI (blue)., Left three images of ducts, right three from alveoli. Scale bars: 15µm., Arrows point to representative SOX9+, Basal (yellow + cyan), Luminal (yellow + white), or Basal/Lum1inal hybrid cells (yellow, cyan + white) based on CK expression. Scale bars: *20µm*., E Quantification of percent SOX9+ Basal, Luminal, or Basal/Luminal hybrid epithelial cells of all epithelial cells per tissue section using the cytokeratin staining combinations denoted in D., F Quantification of percent SOX9+ Basal/Lum1inal hybrid epithelial cells of all Basal/Luminal hybrid epithelial cells.. Error bars represent the mean ± SD (Nullip n = 5, Lac n = 4, 2 wks n = 5, 1 m10 n = 4, 3 mo n =5, 4-6 mo n = 4).. Ordinary one-way ANOVA Sidak’s Multiple comparisons test. No asterisk p > 0.05, *\*p* ≤ 0.05.

SEMA7A+ cells were also increased within the breast epithelium following weaning, which peaked at 1 month post-wean and decreased by 4-6 months post-wean (**Fig. 5A+B**). The percentages of SEMA7A high-expressing cells were most abundant in nulliparous tissue and at 1 month post-wean, similar to what was observed with SOX9+ cells, but was not maintained in the late post-wean samples (**Fig. 5C**). Approximately 40-60% of SEMA7A+ cells were classified as luminal, 15-30% as basal, and 20-30% as basal/luminal hybrid cells in most samples with the exception of the 2-week timepoint samples where SEMA7A+ epithelial cells were approximately 15% luminal, 70% basal, and 15% luminal or basal/luminal hybrid cells (**Fig. 5E**). Further, SEMA7A+ basal/luminal hybrid cells coincided with peak SEMA7A levels observed in all epithelial cells at 1 month post-wean (**Fig. 5F**).

**Fig. 5.**
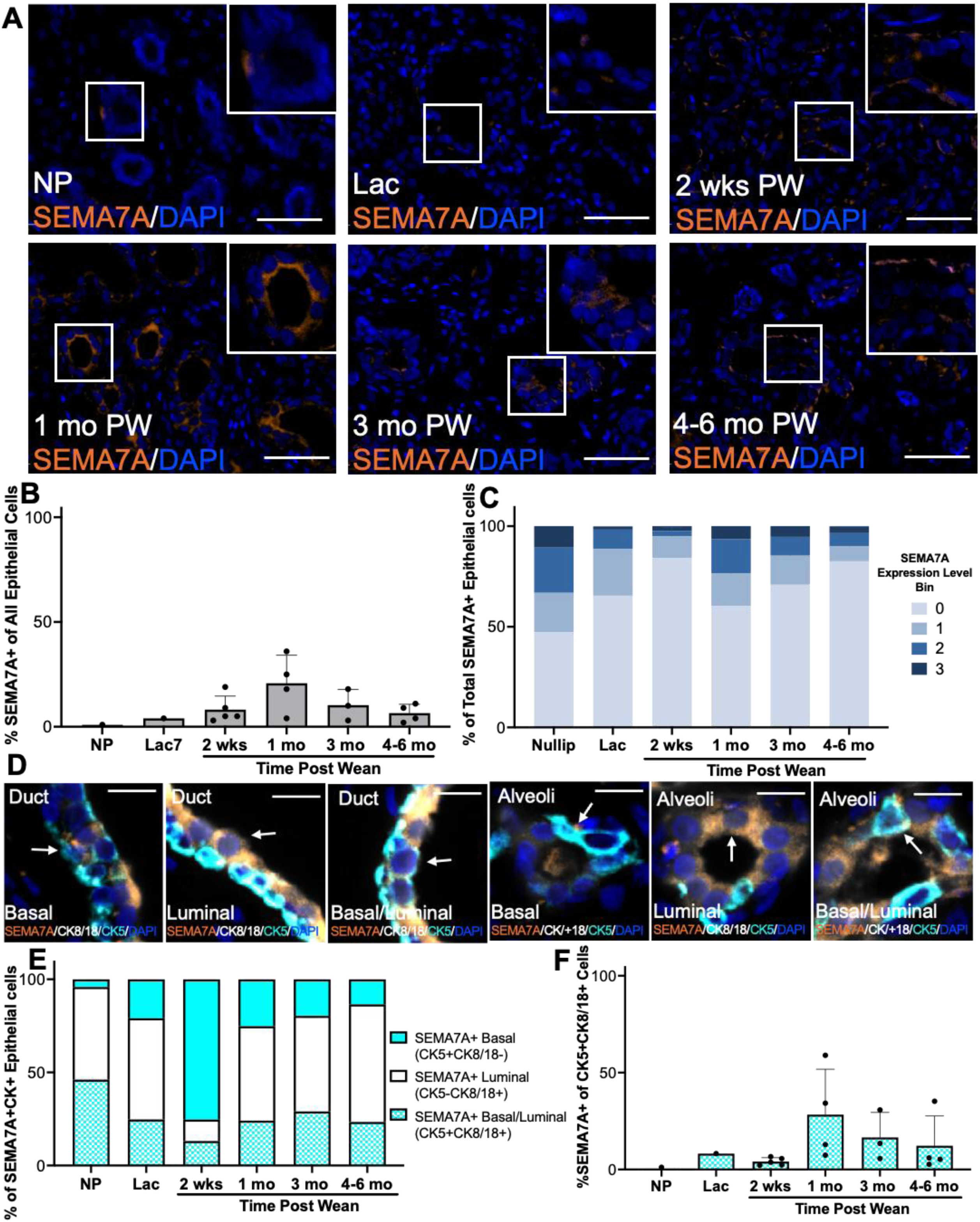
SEMA7A expression in the human mammary epithellium peaks at 1 mo post-wean. **A** Representative multi-spectral IHC (mlHC) image of human mammary tissue stained for SEMA7A (orange) at indicated Nulliparous (NP), Lactation (Lac), and Post-Wean (PW, Involution) time points; nuclei stained with DAPI (blue).. Scale bars: 50 *µ* m. **B** Quantification of percent SEMA7A+ epithelial cells of all epithelial cells per tissue section. **C** Quantification and binning of percent SEMA7A expression level from1low (0) to hig1h (5) of total SE.MA7A+ epithelial cells per tissue section. **D** Representative mlHC images of human mammary tissue stained for SEMA7A (orange), Cytokeratin5 (CK5 cyan), and Cytokeratin 8/18 (CK8+18 white), showing SEMA7A+ CK5+CK8/18-Basal cells, CK5-CK8/18+ Luminal cells, and CK5+CK8/18+ Basal/Luminal hybrid cells; nuclei stained with DAPI (blue). Left three images of ducts, right three from1alveoli.. Scale bars: 15µm. Arrows point to representative SEMA7A+, Basal (orange + cyan), Luminal (orange + white), or Basal/Luminal hybrid cells (orange, cyan + white) based on CK expression.. Scale bars: *20µm..* **E** Quantification of percent SEMA7A+ Basal, Lum1inal, or Basal/Luminal hybrid epithelial cells of all epithelial cells per tissue section using the cytokeratin staining1 combinations denoted in D. **F** Quantification of percent SEMA7A+ Basal/Luminal hybrid epithelial cells of all Basal/Luminal hybrid epithelial cells. **B+F** Error bars represent the mean ± SD (NP n = 1, Lac n = 1, 2 wks n = 5, 1 mo n = 4, 3 mo n =3, 4-6 mo n = 4). Ordinary one-way ANOVA Sidak’s Multiple comparisons test. No asterisk p > 0.05..

### SOX9 and SEMA7A are co-expressed in breast cells and in tumors

To examine whether SOX9 and SEMA7A are co-expressed in the breast epithelium, dual positivity was assigned when both SOX9 and SEMA7A fluorescent signals were detected within the same segmented cell (**Fig. 6A**). A subset of SOX9 and SEMA7A co-expressing cells also appeared to be luminal/basal hybrid cells based on co-expression with CK5 and CK8/18 (**Fig. 6B**). In addition, the percentage of epithelial cells exhibiting dual positivity was increased in the epithelium of samples collected 1 month post-wean while SOX9-SEMA7A co-expressing cells were rare in the other samples (**Fig. 6C**). To determine whether SOX9 and SEMA7A are co-expressed in human epithelial tissue more broadly, we analyzed a publicly available scRNA-seq dataset from normal human breast tissue for *SOX9* and *SEMA7A* mRNA expression^105,106^. Consistent with our scRNA-seq analysis of mouse mammary tissue and our evaluation of human mammary tissues, *SOX9* mRNA was more abundant than *SEMA7A* across MEC populations, consisting of luminal adaptive secretory precursor cells (LASPs), basal-myoepithelial cells, and luminal hormone-sensing cells (**Supplemental Fig. 15A+B**). Given the similarities between LASPs and LPCs^105,107^, we next examined the relationship between *SOX9* and *SEMA7A* in LASP transitional regions that overlapped with other cell populations, as suggestive of adaptive or hybrid cell states. We first narrowed in on *SEMA7A*-expressing cells, since *SEMA7A* was less abundant, and subsequently assessed *SOX9* expression within those SEMA7A-enriched regions. We identified *SOX9* and *SEMA7A* expression in LASP regions that overlapped with basal myoepithelial cell populations, and within a mixed region consisting of LASPs with basal-myoepithelial, luminal hormone-sensing cells, and immune cells (**Fig. 6D**).

**Fig. 6.**
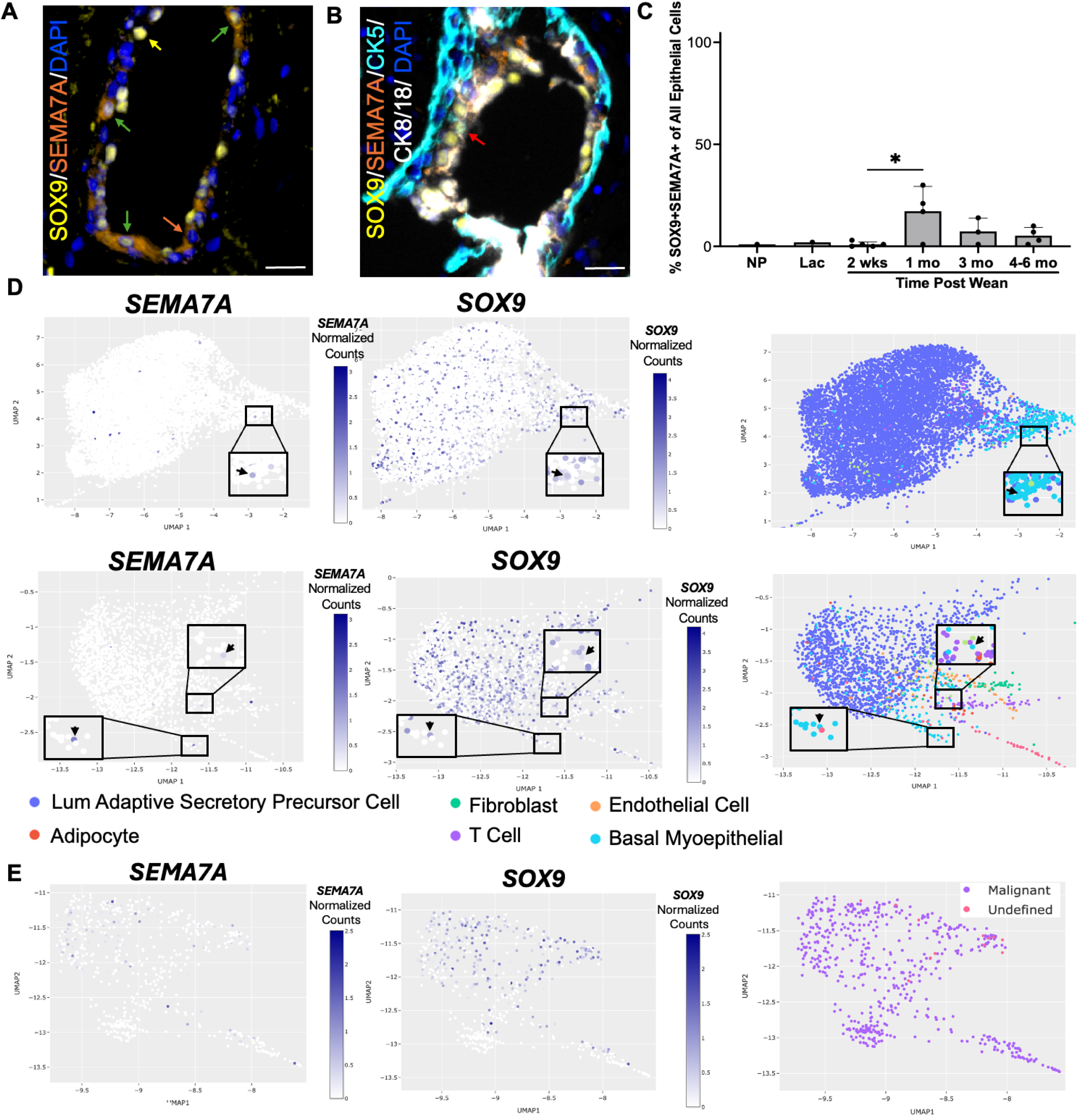
*SOX9* and *SEMA7A* are co-expressed in breast epithelium during involution and in breast tumors. **A** Representative image of human mammary tissue stained for SOX9 (yellow) and SEMA7A (orange) cells; nuclei stained with DAPI (Blue). Arrows denote representative SOX9+SEMA7 A-(yellow arrow), SOX9-SEMA7 A+ (orange arrow), and SOX9+SEMA7 A+ coexistence (green arrow). **B** Representative image of human mammary tissue stained for SOX9 (yellow), SEMA7A (orange), CK5 (cyan), and CK8/18 (white); nuclei stained with DAPI (blue). Red arrow denotes representative SOX9+SEMA7A+CK5+CK8/18+ luminal/basal hybrid cell Scale bars = 20 *µ* m. C Quantification of percent dual SOX9+SEMA7A+ epithelial cells of all epithelial cells. **D** Close-up of Luminal Adaptive Secretor Precursor cells (LASPs) clusters from scRNA-seq analysis of normal human breast tissue overlayed with normalized counts of *SEMA* 7*A* (left) or *SOX9* (middle) mRNA expression, shown compared to annotated cluster (right). Boxes denote regions of *SEMA7A* and *SOX9* co-expression. From Bhat-Nakshatri et. al, 2024 data set, n = 66 samples. **E** Close-up of malignant cluster overlayed with normalized counts of *SEMA7A* (left) or *SOX9* (middle) mRNA expression shown compared to annotated cluster (right). From Qian et. al, 2020, shRNA-seq analysis of human breast tumors. n = 7 samples.

Ulted in breast cancer progression, we hypothesized that co-expression may also occur in breast tumors and promote progression. While the lack of parity status precludes our ability to obtain data on PPBC in publicly available mRNA datasets, analysis of scRNA-seq data from human breast tumors revealed that both *SOX9* and *SEMA7A* mRNA expression is abundant in specific regions of malignant and undefined breast cells^108^ (**Fig. 6E; Supplemental Fig. 15C+D**). We next queried for *SOX9* mRNA expression in The Cancer Genome Atlas breast cancer (TCGA-BRCA) data set, where we have previously shown that *SEMA7A* expression is elevated in all subtypes compared to normal^66,109^. In contrast to *SEMA7A*, *SOX9* was decreased in both luminal and human epidermal growth factor receptor 2-positive (HER2+) tumors; however, expression was increased in TNBC tumors compared to normal tissue (**Fig. 7A**). Next, we evaluated co-expression of *SEMA7A and SOX9* mRNA in the major TNBC subtypes within this breast cancer dataset and observed that, in comparison to normal, *SOX9* and *SEMA7A* are both significantly upregulated in the mesenchymal subtype (M), which is enriched for genes relating to cell motility, TGF-*β* signaling, and Wnt/*β*-catenin signaling, and the unstable subtype (UNS), which is classified as genomically unstable, and is strongly correlated with a basal-like molecular classification^110^(**Fig. 7B**). Consistent with this, *SOX9* and *SEMA7A* are also upregulated in the basal-like 2 (BL2) subtype, although this did not reach statistical significance. In an additional dataset, the Sweden Cancerome Analysis Network – Breast (SCAN-B), of early-stage breast cancers, analysis of *SOX9* and *SEMA7A* confirmed that both are significantly increased in TNBC primary tumors compared to non-TNBC tumors (**Fig. 7C**). Having published that *SEMA7A* expression promotes decreased 5-year overall survival in the SCAN-B dataset^78^, we also observe a similar effect with elevated SOX9 (**Fig. 7D**). Next, Kaplan-Meier analysis was performed to examine the association between *SEMA7A and SOX9* co-expression and DMFS in all breast cancer patients since *SEMA7A* was upregulated in all subtypes. We observed that high *SEMA7A* expression only significantly decreased DMFS in tumors with high *SOX9* expression at both 5-year and 10-year follow up periods (**Fig. 7E, Supplemental Fig. 15E**). Further, stratification of patients by tumor ER status revealed that *SEMA7A* expression in *SOX9* high tumors was associated with reduced DMFS in both ER- and ER+ subsets, supporting the idea that this dyad promotes metastasis independent of ER status (**Fig. 7F**). Additionally, patients with ER-tumors demonstrate a greater reduction in DMFS within the 5-year follow-up period, whereas patients with ER+ tumors exhibited a more pronounced decrease over 10-years, consistent with prior literature showing that ER+ breast cancers tend to recur later than ER- or TNBC tumors^111,112^.

**Fig. 7.**
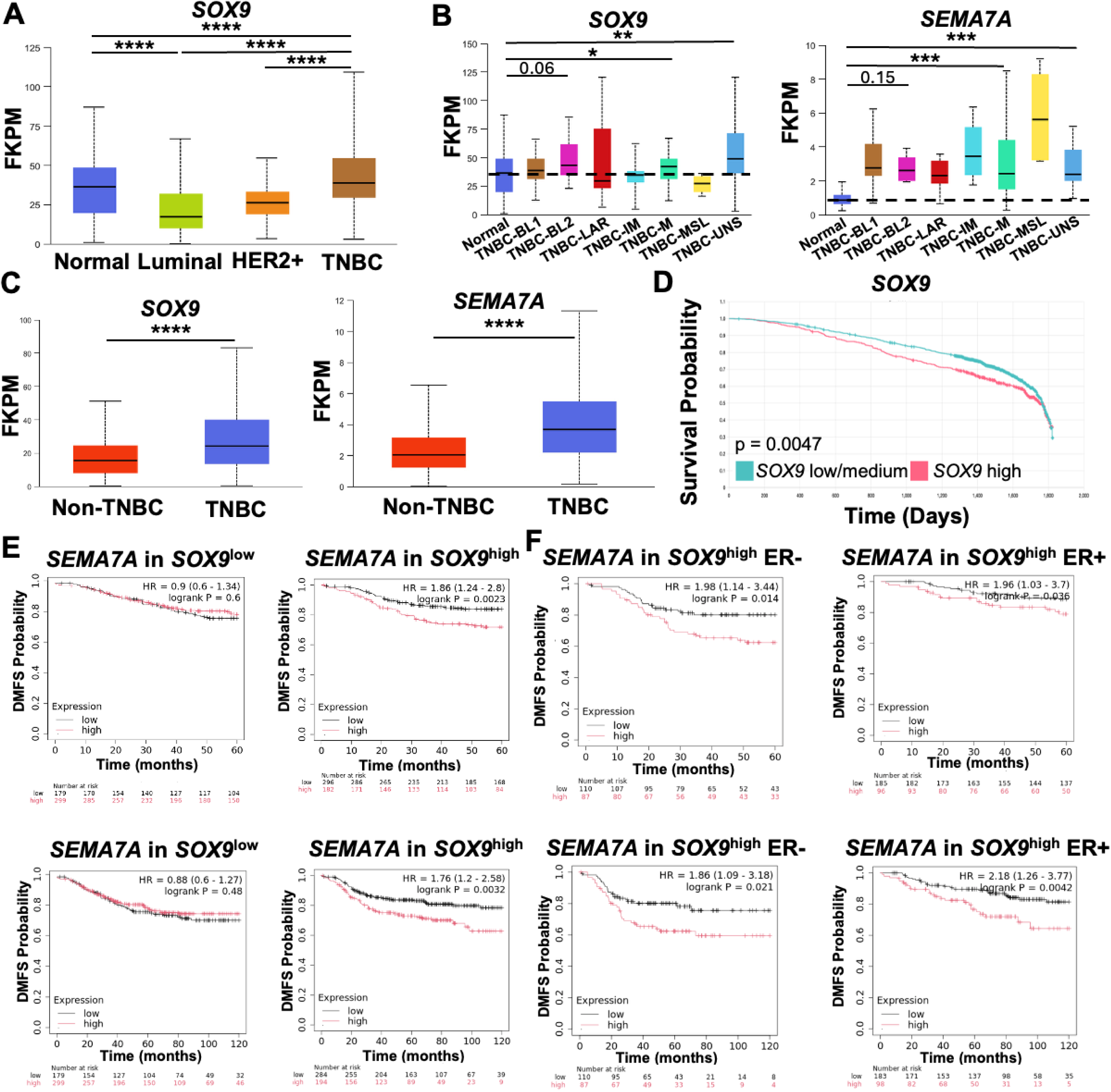
SOX9 and SEMA7A co-expression is associated with poor outcomes in breast cancer. **A** *SOX9* mRNA expression in normal mammary tissue versus breast cancer primary tumor samples separated by major breast cancer subtypes (Luminal = 575, HER2-positive n = 37, TNBC n = 115), from TCGA. **B** *SOX9* (left) or *SEMA7A* (right) mRNA expression in breast cancer primary tumor samples separated by TNBC subtypes (Basal-Like 1(BL-1) n = 13, Basal-Like 2(BL-2) n = 9, Luminal Androgen Receptor (LAR) n = 8, lmmunomodulatory (IM) n = 18, Mesenchymal (M) n = 27, Mesenchymal Stem-Like (MSL) n = 4, Unstable 1(UNS) n = 20), from TCGA. Dotted line denotes median expression level in normal breast tissue. C *SOX9* (left) or *SEMA7A* (right) mRNA expression (FPKM) in Non-TNBC versus TNBC primary tumor samples (Non-TNBC n = 8332, TNBC n = 874), from SCAN-B. D Breast cancer patient Kaplan-Meier 2000-day overall survival analysis of low/medium and high *SOX9* mRNA expression (low/medium n = 1935, high n = 645). E Kaplan-Meier 5- (top, 60-month) and 10-year (bottom, 120-month) distant metastasis free survival (DMFS) analysis of low and high *SEMA7A* mRNA expression in SOX9 low mRNA expressing (left) or s,OX9 high mRNA expressing tumor samples (right) from all subtypes of breast cancer combined. F Kaplan-Meier 5-year (top, 60-month) and 10-year (bottom, 120-month) DMFS analysis of low and high *SEMA7A* mRNA expression in SOX9 high mRNA expressing tumor samples separated by ER- (left) or IER+ (right) tumors. E+F From KM-Plotter. A-C Box plots are median (black line), ± upper and lower quartiles (colored box), and maximum and minimum values (whiskers). For C, significance shows subtypes where *both SOX9* and *SEMA7A* expression are significantly increased compared to normal. *\*p* ≤ 0.05, *\*\*p* ≤ 0.01, *\*\*\*p* ≤ 0.001. *\*\*\*\*p* ≤ 0.0001.

## DISCUSSION

The mammary epithelium remains functionally plastic throughout adult life in female mammals, and full differentiation is not achieved until milk secretion during lactation. The necessity for cellular reprogramming to adapt to complex changes that occur to support lactation and those that must occur post-lactation – during postpartum involution – renders the tissue susceptible to oncogenic transformation. Postpartum mammary involution has proven a tumor-promotional developmental window by multiple studies showing durable alterations to protein expression and pro-oncogenic phenotypes by tumor cells exposed to this developmental event. Here, we identify novel co-expression of SOX9 and SEMA7A in mammary LPCs during postpartum involution in murine mammary glands, validate our findings in a unique cohort of human post-lactational breast tissues, and reveal that co-expression is co-opted by breast cancers and promotes poor prognosis in both ER+ and ER-subtypes.

Using scRNA-seq of mouse MECs from actively lactating, or those undergoing involution, we reveal that *Sox9* expression is high in both luminal and basal myoepithelial populations and is enriched in LPCs. We characterize for the first time SOX9 protein expression in post-lactational murine mammary and human breast tissues and show that SOX9 is highly expressed in nulliparous tissues, decreases in lactation samples, and is reestablished to nulliparous levels during weaning-induced involution. We also observed a small subset of LPCs express *Sema7a*, and that *Itgb1* and *Itga6*, key SEMA7A receptors^66^, are expressed more widely. Of the semaphorin family of proteins, aptly named based on their ability to send messages from one cell to another, SEMA7A is the only GPI-membrane-linked member^113^, and it can also be shed to signal in an autocrine manner^76^. This suggests that a small population of SEMA7A-expressing cells has the capacity to influence neighboring cells that are poised to respond to SEMA7A through expression of integrin receptors. This could account for SEMA7A having broader implications in the survival of LPCs during involution, which is supported by our previous observation that live LPCs are significantly decreased during involution in SEMA7A KO mice, resulting in accelerated cell death and tissue remodeling^66^. A key limitation of our study was our focus on EpCAM+CD45-cells to identify MEC subsets, which are typically characterized by their expression of EpCAM and CD49f. However, our *in vitro* studies in cultured MECs identify a highly mesenchymal subset of cells that are EpCAM-CD45f-that are driven by forced SEMA7A expression. These cells would often be classified as stromal cells in the mammary epithelium, which are not present in cultured MECs, and suggest an active role for mesenchymal reprogramming by SEMA7A in epithelial cells. Accordingly, our future studies will be guided by this observation and will expand the single-cell analysis to include in-depth pathway analysis in EpCAM-populations and other cell types expressing SEMA7A from our current analysis (**Supplemental Fig. 4**).

Furthermore, our dataset also contains EpCAM-CD45+ immune populations, which future studies will investigate to provide additional insight into SEMA7A during involution beyond LPCs.

Building on our *in vivo* observations, consequences of decreased *Sox9* in MEC models of lactation and involution (LadDiff/Inv) include impaired lactogenic differentiation, increased mesenchymal cell phenotypes, and increased SEMA7A expression; this may suggest that SOX9 acts as a transcription repressor of SEMA7A in MECs to maintain epithelial cell identity. SEMA7A may also mark a subset of highly mesenchymal mammary cells that resemble mesenchymal stem-like cells, which is supported by bulk RNA-seq showing enrichment for EMT-associated transcriptional programs when SEMA7A is expressed. In addition, activation of distinct programs with *Sox9* KD and S7OE suggests that the convergence on a mesenchymal cell state may occur through overlapping and mechanistically distinct transcriptional programs. In support of this, *Sox9* and *Sema7a* mRNA are largely expressed in distinct LPC populations, and our data raise the possibility that *Sema7a* expression contributes to the acquisition of mesenchymal-like phenotypes in LPCs, which drive LPCs toward basal/myoepithelial-like states that are proposed as the cells of origin for basal-like breast cancers^114^. Finally, both *SOX9* and *SEMA7A* expression are increased in the TNBC-Mesenchymal (TNBC-M) and unstable (UNS) subtypes of TNBC, which are associated with the basal-like breast tumor classification^110,115^. Therefore, co-expression of SOX9 and SEMA7A, particularly in LPC populations, may create a cellular context especially conducive to malignant transformation.

Our studies also identified a transient increase in the dual SOX9+SEMA7A+ epithelial populations in human breast tissues in early post-wean samples, highlighting a window of coordinated pathway activation during involution. Notably, prior analysis of this same tissue cohort demonstrated that at 1-month post-wean, there is an equal distribution of lobule types, and dilated and collapsed lymphatic vessels that correlate with cyclooxygenase-2 (COX-2) expression^96^, known for its activity in involution-associated tumor growth, lymphatic remodeling, invasion, and metastasis^13,15,16^. We have also shown that SEMA7A expression is dependent on COX-2 expression^82^ and both are sufficient for tumor cell invasive phenotypes, lymphangiogenesis, and metastasis^77,82^. Furthermore, SOX9+ or SEMA7A+ epithelial cells displayed hybrid epithelial phenotypes, and scRNA-seq from normal human breast tissue^105^ revealed *SOX9* and *SEMA7A* co-expression within LASP populations that exhibit increased lineage plasticity. Therefore, our findings, when considered in the context of published studies, identify the 1-month post-wean period as a transient window during which SOX9 and SEMA7A are co-expressed in normal MECs, whereby sustained co-expression may result in transformed MECs that also contribute to a pro-lymphangiogenic environment to facilitate metastatic spread. Collectively, these results support that this transient window of heightened co-expression during involution may represent a therapeutic window for targeted intervention, as exemplified by our studies using COX2 inhibitors to prevent PPBC metastasis^16,116,117^.

Historically, data on parity status, such as time since last childbirth, which defines PPBC, have not been extensively collected. However, in general breast cancer datasets that do not account for parity status, we were able to show that both *SOX9* and *SEMA7A* are elevated in TNBC tumors compared to normal tissue, and that high co-expression confers increased risk for metastasis across all breast cancers regardless of ER expression. Aligning with the more aggressive nature of TNBC, which recurs early, co-expression in ER-tumors was associated with a more pronounced reduction in DMFS within the first five-years of follow-up. In contrast, in ER+ breast cancers, which tend to recur later, co-expression confers significantly increased risk for metastasis at the 10-year follow-up period, highlighting a role for the SOX9-SEMA7A axis in late-stage recurrence, a major challenge in ER+ disease^111,112^. This indicates that while SOX9 and SEMA7A may initially play a larger role in promoting metastasis in ER-tumors, they may also contribute to disease progression and recurrence in late-stage ER+ tumors, potentially through adaptive resistance to ER-targeted therapies, as supported by our previous studies of SEMA7A^83,85^ and studies from others on SOX9 in ER+BC^118^.

Collectively, our results suggest that SOX9 and SEMA7A are co-expressed on distinct cell populations in the postpartum mammary gland to regulate cellular plasticity during involution in mouse and human tissues, and this co-expression may be co-opted by breast tumors to drive alterations in cellular plasticity and poor prognosis. Acquisition of a SOX9+SMEA7A+ phenotype may become more robust with successive pregnancies, as suggested by our studies showing SOX9 expression in MECs after involution and by mouse studies in which parity induces lasting alterations to the mammary transcriptome that shape cellular responses during subsequent rounds of pregnancy, lactation, and involution^33,54–56^. Further, SOX9 and SEMA7A co-expression emerges in our study as a potential driver of breast cancer progression more broadly, revealing an axis that may promote metastasis across all breast cancer subtypes. In related studies, we have developed and tested a novel monoclonal antibody against SEMA7A in mouse models that decreases tumor growth, tumor-associated lymphatic vessels, immunosuppression, and prevents metastatic outgrowth in multiple models of TNBC and ER+ breast cancer with no evidence of toxicity^12,78,84,85^. Therefore, targeting SEMA7A represents a novel therapeutic intervention that could be applied both as a preventative in the postpartum setting, as well as in the treatment of PPBC and all aggressive breast cancer subtypes where SEMA7A expression is observed, which has the potential to impact thousands of breast cancer patients.

## RESOURCE AVAILABILITY

### Lead contact

Further information and requests for resources and reagents should be directed to and will be fulfilled by the lead contact, Traci Lyons, PhD.

### Materials availability

This study did not generate new, unique reagents.

### Data and code availability

- This paper analyzed publicly available human datasets
- Single-cell RNA-seq data have been deposited at GEO at GSE313223 and are publicly available as of the date of publication.
- Bulk RNA-seq data have been deposited at GEO at GSE319717 and are publicly available as of the date of publication.
- This paper does not report original code.
- Any additional information required to reanalyze the data reported in this paper is available from the lead contact upon request.

## METHODS

### Single Cell RNA-sequencing

All animal experiments were approved by the University of Colorado (CU) Anschutz Medical Campus Institutional Animal Care and Use Committee. Female C57BL6/J mice were bred with males to induce pregnancy. Right and left #4 mammary glands were isolated from 11-week-old (at time of harvest) females, either after 10 days post-parturition (Lac10), or after 10 days of lactation, followed by removing pups to force weaning and initiate involution for 3 days (Inv3). Inguinal lymph nodes were removed, and the mammary glands were digested to single-cell suspensions using protocols previously described^66^. Single Cells were resuspended in 1% UltraPure non-acetylated BSA (Invitrogen #AM2616) in Magnesium and Chlorine-free PBS, and stained with anti-EpCAM antibody (APC-Cy7, 1:200, BioLegend, # 118218) or anti-CD45 antibody (FITC, 1:200, BioLegend #103107) for 30 minutes at 4°C. Cells were washed with 1% BSA-PBS and resuspended in 1% BSA-PBS containing a 1:1000 dilution of 7-AAD (Thermofisher Scientific: #A1310) for dead cell exclusion. UltraComp eBeads Plus Compensation Beads (Invitrogen, #01-3333-42), fluorescence minus one (FMO) stained cells, and unstained cells were also used as staining controls. Immediately, cells were run on a MoFlo XDP Flex cell sorter (Beckman Coulter) with a 100um nozzle, in conjunction with the CU Flow Cytometry Shared Resource. Debris, doublets, and dead cells were excluded, and FACS gates were set based on single stain, FMO, and unstained controls (**Supplementary Fig. 1B**). Equal proportions of EpCAM+CD45- and CD45+ cells were sorted and pooled (70,000 of each cell type for a total of 140,000 cells) into 1% BSA-PBS buffer. Post-sort, cells were spun down (200rcf for 5 minutes) and resuspended in 0.1% BSA-PBS. Single-Cell 3’ Gene Expression profile was obtained using the Chromium GEM-X Single-Cell 3’ Library and Gel Bead Kit (10x Genomics). After cell counting and viability assessment, 20,000 viable flow-sorted cells were loaded on the Chromium X instrument (10x Genomics) for single-cell capture, followed by amplification of cDNA and library preparation to recover 15,000 cells per sample. Libraries were quantified, pooled, and sequenced on the NovaSeq X plus sequencer (Illumina) at 50,000 reads/cell at the CU Genomics Shared Resource.

Raw base call files were demultiplexed and FASTQ sequencing files were produced by the CU Genomics Shared Resource. FASTQs were processed with Cell Ranger 9 (10x Genomics) on a SLURM HPC cluster. Per-sample quantification was performed using the ‘cellranger9 count’ was run for each library with default chemistry autodetection and using the reference 10x mouse transcriptome refdata-gex-mm10-2020-A. Libraries were aggregated using ‘cellranger9 aggr’ to generate the final filtered, raw feature-barcode matrices and Loupe Browser explorer files. Downstream analysis was performed in R 4.4.2 and rendered with Quarto 1.8.25. Count matrices were imported, and ambient RNA was estimated and corrected with SoupX 1.6.2. Corrected counts were used to create per-sample Seurat 5.3.0 objects. Quality control filtering and doublet detection used scuttle 1.16.0 and scDblFinder 1.20.2, respectively. After filtering, samples were merged into a single Seurat object for normalization, dimensionality reduction, clustering, and cell type annotation. Annotations were determined by manual review of canonical markers, module scores for curated gene sets, and automated reference matching with clustifyr 1.18.0, followed by final curation in a cluster-to-label mapping file. To further characterize specific cell populations of interest (e.g., epithelial and stromal subsets), we subsetted the data to include only relevant clusters (Alveolar Luminal, Luminal Progenitor, Luminal Mature, Basal/Myoepithelial, and Pericytes). These subsets were then re-processed, starting from normalization: variable features were re-identified, data were scaled, and PCA was recalculated. We then performed sub-clustering (resolution = 0.5) and generated new UMAP visualizations to resolve finer-scale heterogeneity within these lineages. Differential expression was performed within each annotated cell type, comparing genotypes pairwise within a condition using the Wilcoxon rank-sum test implemented in Seurat::FindMarkers. Genes were considered for testing if they were expressed in at least 25% of cells in either of the two groups being compared (min.pct = 0.25). Significance was defined based on an adjusted p-value (Benjamini-Hochberg correction) of < 0.05 and an absolute log2 fold-change threshold of > 0.25. Pathway enrichment analyses were conducted on significant genes with clusterProfiler 4.14.6 using MSigDB gene sets via msigdbr 24.1.0 and mouse annotations from org.Mm.eg.db v3.20.0. Loupe Browser v9.0.0 (10X Genomics) was used to generate the plots in **Fig. 2A**.

### Immunohistochemistry

Formalin-fixed and paraffin-embedded C57BL/6 female mouse mammary tissues were utilized from previously described studies^66^. Antigen retrieval was performed as previously described for CC3^66^, E-Cadherin, Vimentin^84^, Ki67^82^, and SOX9 (1:100, Cell Signaling Technologies, #82630S). For secondary staining, the EnVision+ Single Reagent HRP.Rabbit (Agilent, #K400311-2) was used and visualized with the ImPACT DAB Substrate (Vector Laboratories, #SK-4015). Hematoxylin was used for the counterstain (Vector Laboratories, #H-34-1-500). Slides were scanned using a 20X objective on an Olympus BC63 microscope with a DP27 camera (Evident Scientific), and images were acquired using cellSense imaging Software (Evident Scientific). FIJI open-source software^119^, was used to convert the output .tif images to pyramidal .tif files for Qupath for functionality reasons. Qupath open-source software (Version 6.0) ^120^was used to create an automated analysis pipeline to quantify SOX9+ cells within the epithelium. Annotations were created on each image, capturing all tissue except for large portions of muscle tissue. The “estimate stain vectors” function in QuPath was used to standardize the SOX9 stain. InstanSeg, an opensource extension for Qupath (Goldsborough arXiv:2408.15954) that detects and segments cells on brightfield images, was used to identify and label cells with the following settings: Preferred device: GPU; Threads: 4; Tile Size: 512; Tile Padding: 32; Input Channels: Red, Green, Blue; Outputs: Only Channel; Output Type: Default; Make Measurements: True; Random Colors: False. The neural network machine learning classifier was used to classify segmented cells as “epithelium” or “stroma” and trained on an arbitrarily selected region from an image for each reproductive time point, with a maximum of 1,000 iterations. This neural network used segmented cell features to classify cells into each category; the features included are listed in **Supplemental Table 1**. To better classify the cells, smoothed object features were added at 25µm and 50µm radius to the existing InstanSeg measurements.

Ground truth annotations were added to the epithelial and stromal categories for the selected training regions until we were satisfied with the neural network’s ability to classify cells into the two categories. A second neural network was created for tissue samples collected during lactation because the histological morphology was distinct from involution and nulliparous time points. Lastly, color thresholding of the SOX9 DAB stain was used to separate each cell into one of 3 bins with the following threshold values, respectively: bin 1: 0, bin 2: 0.2, bin 3: 0.4. An automated script pipeline was created from all these steps and then applied to each annotation for each respective image. Final epithelial cell amounts from each annotation were then summed, and the totals from bins 2 and 3 (SOX9high) were used to calculate the percentage of SOX9+ cells in the epithelial tissue.

### Cell Culture

Culture and differentiation of murine HC-11 and EpH4 mammary epithelial cell line methods were adapted from Geletue et al. ^88^. HC-11 cells were obtained from Weston Porter, PhD (Texas A&M University). HC-11 cells were maintained in RPMI 1640 with 2.05 *m*M L-glutamine (Cytiva HyClone, #SH30027.01), 10% FBS (Peak Serum, #PS-FB3), 5 *u*g/mL human-recombinant Insulin (Gibco, #12585-014), and 10 *n*g/mL human-recombinant EGF (Corning, #354052). EpH4 cells were obtained from Stephen Anderson, PhD (CU Anschutz Medical Campus) and were maintained in the same media as HC-11 cells but without EGF. To induce lactogenic differentiation (LacDiff), HC-11 or EpH4 cells were grown to confluence in standard maintenance media, and 24 hours later, the maintenance media was replaced with priming media containing RPMI 1640 with 2.05 *m*M L-glutamine, 10% FBS, and 5 *u*g/mL human-recombinant Insulin, for which cells were maintained for an additional 24 hours. Priming media was then replaced with differentiation media consisting of RPMI 1640 with 2.05 *m*M L-glutamine, 10% FBS, 1 *u*g/mL Hydrocortisone (Sigma, #H4001), 5 *u*g/mL human-recombinant Insulin, and 100 *n*g/mL human-recombinant Prolactin, (R&D Systems, #628-PL, reconstituted in Reconstitution Buffer 4 (BSA/HCL), R&D Systems, #RB04). Differentiation media was replaced every other day for 7 days, and cultures were monitored for dome formation. On day 7 of differentiation, when dome formation and milk protein production are high, the differentiation media was swapped for priming media, mimicking hormone withdrawal and involution (Inv). Cells were maintained in priming/involution media for 4 days, swapping media on day 2, to allow for sufficient cessation of milk protein production and an increase in apoptosis. Cells continuously cultured in priming media that never received lactogenic hormones were used as a control (Ctrl). Cell lines were regularly tested for mycoplasma (Lonza, #LT07). All brightfield images of cells were obtained on a Zeiss Axio Vert.A1 microscope with an AxioCam MRc camera (Zeiss) using Zeiss Zen software.

### shRNA knockdown and overexpression

To generate SOX9 KD cell lines, plasmids containing SOX9 shRNA constructs (The RNAi Consortium (TCR) shRNA Library, Broad Institute, pLKO.1-puro backbone, KD: Clone #TCRN0000086165), packaged into lentivirus, was obtained from the University of Colorado Functional Genomics Shared Resource. The same backbone vector containing a non-targeting shRNA was used as a control (Sigma Aldrich, #SHC016). To generate SEMA7A OE cell lines, plasmids containing a SEMA7A-ORF Clone insert (Origene, #MR209876L3, pLenti-C-Myc-DDK-P2A-puro backbone) were packaged into lentivirus at the CU Functional Genomics Shared Resource. The same backbone without an ORF insert was used as an empty vector control (Origene, #PS100092). HC-11 and EpH4 cells were grown to 70% confluence and incubated with a 1:1 ratio of cell maintenance media and viral particles with 8 *u*g/mL polybrene (Sigma-Aldrich, #TR-1003). 48 hours later virus was removed, and cells were allowed to recover in complete maintenance media for 24 hours. Transduced cells were then selected for puromycin resistance (5 *u*g/mL, Gibco, #A113803). SOX9 KD or SEMA7A OE was regularly confirmed by qPCR and western blot in undifferentiated cells and at various experiment endpoints.

### Bulk RNA-sequencing

HC-11 and EpH4 shCtrl, shSOX9, EV, and S7OE cell lines, grown in 3 separate dishes per cell line to generate technical triplicates, were subject to our lactogenic differentiation protocol, as described. On involution day 4, cells were lifted with Accutase (Gibco, #A1110501), and 200,000 cells per sample were re-suspended in 50ul of DNA/RNA Shield buffer (Zymo Research, #R1100-50) and sent to Plasmidsaurus for sequencing. Total RNA extraction and sequencing were performed by Plasmidsaurus using Illumina sequencing and 3’ end counting, with custom analysis and annotation. Prior to sequencing, total RNA was extracted from cells using a bead-based extraction approach, and RNA sample concentrations were calculated and normalized. For sequencing library preparation, mRNA was converted into cDNA via reverse transcription and second-strand synthesis, followed by segmentation, library indexing, and amplification.

Analysis was also performed by Plasmidsaurus, in which FASTQ sequence files were generated, and demultiplexing was performed with BCL Covert v4.3.6 and fqtk v0.3.1. Read-filtering using FastP v0.24.0 was performed, including poly-x tail tripping, 3’ quality-based tail trimming, a minimum Phred quality score of 15, and a minimum length requirement of 50bp. Alignment to the reference genome was performed using STAR aligner v2.7.11 with non-canonical splice junction removal and output of unmapped reads. Coordinate sorting of BAM files was then performed using samtools v1.22.1. For UMI-based de-duplication, removal of PCR and optical duplicates was conducted using UMICollapse v1.1.0. For mapping QC, alignment quality metrics, strand specificity, and read distribution across genomic features were evaluated using RSeQC v5.0.4 and Qualimap v2.3, and a QC report was generated using MultiQC v1.32. Gene-expression was quantified using feature Counts (subread package v2.1.1) with strand-specific counting, multi-mapping read fractional assignment, exons and 3’ UTR as the feature identifiers, and grouped by gene_id. Final gene counts were annotated with gene biotype and other metadata extracted from the reference GTF file. Sample-sample correlations for sample-sample heatmaps were calculated on normalized counts. Differential expression was determined using edgeR v4.0.16 with filtering for low-expressed genes with edgeR::filterByExpr with default values. Finally, functional enrichment pathway analysis was performed using gene set enrichment analysis with GSEApy v0.12 using the MSigDB Hallmark gene set. All graphics displayed for visualization of data were generated using Plasmidsaurus’s provided online analysis workspace.

### Immunofluorescence and Phalloidin Staining

Cells were grown on cell culture-treated coverslips (Neuvitro, GG-18-1.5-PLL) in culture dishes and harvested at various LacDiff/Inv or Ctrl (no lactogenic hormone treatment) time points. At the time of harvest, culture media was removed and cells on coverslips were briefly washed with PBS and fixed in 2% Formalin (Fisher HealthCare, #254685) in PBS for 20 minutes at room temperature. Coverslips were briefly washed with PBS and stored in filter-sterilized 0.1% BSA (Fisher Bioreagents, #BP1600) in PBS at 4°C for short-term storage. Cells were permeabilized with 0.1% Triton-X100 (Fisher Scientific, #BP151) for 15 minutes at room temp and again washed with PBS. Cells were then blocked in 1% BSA-PBS containing 0.1% Tween20 (BSA-PBS-T, BioRad, #1706531) for 1 hour at room temperature, with gentle rocking. To visualize Whey Acidic Protein (WAP) expression, coverslips were incubated with primary antibodies (1:500, Santa Cruz Biotechnology, #sc-398276, Clone E-8) in 1% BSA-PBS-T overnight at 4°C, with gentle rocking. For no primary control samples, coverslips were incubated in 1% BSA-PBS-T without antibodies. The next day, the primary antibody solution was removed, and the coverslips were washed with PBS-T, 3 times for 5 minutes each with gentle rocking. All sample and control coverslips were then incubated with secondary antibodies (FITC, 1:1000, abcam, #ab6785) in 1% BSA-PBS-T for 1 hour, with gentle rocking. Secondary antibody solution was removed, and coverslips were again washed 3×5min in PBS-T. Coverslips were mounted on microscope slides with DAPI-containing aqueous mounting media (abcam, #AB104139). For Phalloidin staining, cells were grown on coverslips, fixed, stored, and permeabilized in the same manner described for IF. Phalloidin (iFluor 488, abcam, #ab176753) was diluted to 1X in 1% BSA-PBS and incubated on coverslips for 1 hour at room temperature. Staining solution was removed, and coverslips were washed with PBS-T, 3×5 minutes with gentle rocking. Coverslips were mounted in the same manner described for IF. All slides were immediately imaged on an Olympus fluorescent microscope IX83 with a Hamamatsu ORCA-Flash 4.0LT camera (#C11440) using a 20X or 40X objective with the same imaging setting across all samples and controls. Pixel size for all images is 0.0069×0.0069 inches^²^. All image display adjustments were conducted in PowerPoint picture corrections and were consistent across all samples and controls (for WAP IF both HC-11+EpH4 cultures: +50% sharpness, +60% brightness, +20% contrast, for Phalloidin: HC-11: +50% brightness, +15% contrast, EpH4:+30% brightness, +15% contrast). For Phalloidin staining, individual cell length (l) and width (w) were analyzed using ImageJ software, and the aspect ratio was calculated using the equation: aspect ratio = width/length.

### Immunoblot

Immunoblots were performed as previously described^84^. Primary antibodies included: SOX9 (1:1000, Cell Signaling Technologies, #82630S), SEMA7A (1:250, Santa Cruz Biotechnology, #sc-374432, Clone C6), E-Cadherin (1:1000, Cell Signaling Technologies, #3195S), Vimentin (1:1000, Cell Signaling Technologies, #5741S, Clone D21H3), and alpha tubulin (1:1000, Cell Signaling Technologies, #9099S, Clone 11H10) as a loading control. Secondary antibodies used included: Goat Anti-Rabbit IgG (1:1000, Cell Signaling Technologies, #7074) and Goat Anti-Mouse IgG (1:1000, Cell Signaling Technologies, #7076).

### Caspase Activity Assays

HC-11 and EpH4 cells were grown in white-walled Costar Assay 96-well plates (Corning, #3903), and a Caspase-Glo 3/7 Assay System (Promega, #G8091) was used according to the manufacturer’s instructions to measure caspase activity in cultures at various stages throughout the *in vitro* LacDiff/Inv process. Luminescence was read on a BioTek Synergy H1M2 plate reader (Agilent).

### qPCR

RNA was extracted from HC-11 and EpH4 cell cultures at the indicated undifferentiated or LacDiff/Inv time points using a Quick-RNA Miniprep Kit (Zymo Research, #R1055) according to the manufacturer’s instructions. RNA from each sample was reverse transcribed to generate cRNA using the iScript cDNA Synthesis Kit (BioRad, #1708890) according to the manufacturer’s instructions. To quantify *Sema7a* (Fwd: CATCGCTCAAAGGCTACCACATG, Rvs: GGTGACTATCAGCTACCTGGAAG), *Sox9* (Fwd: CACACGTCAAGCGACCCATGAA, Rvs: TCTTCTCGCTCTCGTTCAGCAG), *Snail* (Fwd: TGTCTGCACGACCTGTGGAAAG, Rvs: CTTCACATCCGAGTGGGTTTGG), *Zeb1* (Fwd: ATTCAGCTACTGTGAGCCCTGC, Rvs: CATTCTGGTCCTCCACAGTGGA), and *Gapdh* (Fwd: AAGGTCATCCCAGAGCTGAA, Rvs: CTGCTTCACCACCTTCTTGA, as a housekeeping gene) mRNA levels, forward and reverse primers (500 *u*M final concentration) were combined with 100 *u*g of cDNA and iTaq Universal SYBR Green Supermix (BioRad, #1725120). Samples were run and quantified on a QuantStudio 3 Real-Time PCR machine (Applied BioSystems, ThermoFisher Scientific) using the following cycle conditions: 98°C for 30 seconds, 40 cycles: 98°C for 15 seconds, 49.9°C (5°C below the lowest primer Tm) for 30 seconds, 72°C for 30 seconds, followed by a melt curve analysis. To calculate mRNA expression, the following equation was used: expression = 2^-(^Δ^Cq)^, where ΔCq = Cq value of the target gene – Cq value of *Gapdh* (housekeeping gene). Expression values were then normalized to the associated EV or shRNA control sample and reported as fold change expression relative to the control.

### Flow Cytometry

For all flow cytometry experiments, HC-11 and EpH4 cells were lifted with Accutase (Gibco, A110501). Cells were stained with the Zombie UD Fixable Viability Kit according to the manufacturer’s protocol (Biolegend, #423108). Staining of extracellular markers was performed as previously described^84^. Extracellular antibodies included: anti-CD49f (1:200, Pacific Blue, Biolegend, #313620) and anti-EpCAM (CD326, 1:300, BUV737, BD Biosciences, #741818). Cells were fixed and permeabilized using the True-Nuclear Transcription Factor Buffer Set (Biolegend, #424401), and intercellular targets were stained according to the manufacturer’s protocol. Intercellular antibodies included: Vimentin (1:300, Alexa Fluor 647, Biolegend, #699308). UltraComp eBeads Plus Compensation Beads (Invitrogen, #01-3333-42), fluorescence minus one (FMO) stained cells, and unstained cells were also used as staining and compensation controls. All samples were analyzed on a ZE5-YETI cytometer (BioRad) at the University of Colorado Cancer Center (UCCC) Flow Cytometry Shared Resource using the Bioligand Everest software and auto compensation. Auto compensation was confirmed, and data were analyzed using FloJo software (BD Biosciences) using a combination of single stain, FMO, and unstained control cells to determine analysis gates using the gating strategy outlined in **Supplementary Fig. 10**.

### Scratch Wound Migration Assay

HC-11 cells were subject to our *in vitro* LacDiff/Inv process, and after 4 days of Involution, cells were lifted with 0.25% Trypsin-EDTA (Cytiva HyClone, #SV30031.01), and 300,000 cells per well were seeded in 24-well plates in priming/involution media. The next day, confluent monolayers were treated with 10 *u*g/mL of Mitomycin C (Sigma-Aldrich, #M5353) for 2 hours, to inhibit proliferation that could be misinterpreted as migration. After which, Mitomycin C-containing media was removed, and cells were washed with PBS. A scratch was made down the center of each well using a sterile p200 pipette tip. Scratches were washed once with PBS, and cells were topped with priming/involution media. Plates were then loaded on a BioTek BioSpas Live Cell Imaging System (Agilent), and images of the scratch were taken every 4 hours for 24 hours. BioTek Gen5 (Agilent) software was used to measure the percent wound closure using the equation: % wound closure = ((initial wound area − final wound area) / final wound area) × 100.

### Data Mining

For analysis of *SEMA7A* and *SOX9* mRNA expression in publicly available scRNA-seq human normal breast tissue^105^ and breast cancer^108^ datasets, the University of Alabama at Birmingham MammOnc-DB^106^ was used to visualize data and generate graphs. MammOnc-DB was also used to generate graphs of SEMA7A and Sox9 protein and mRNA expression in samples from The Cancer Genome Atlas Breast Cancer dataset (TCGA-BRCA, https://www.cancer.gov/tcga.) and the Sweden Cancerome Analysis Network – Breast^121^, and the SOX9 5-year survival analysis from the SCAN-B dataset. For additional survival analysis, Kaplan-Meier Plotter Breast Cancer gene chip was used^122^ to generate DMFS curves for SEMA7A (Affy ID: 230345_at) and SOX9 (Affy IDL 202936_s_at). Default parameters were used, and percentiles were used to auto-select the best cut-off for dividing patents into high and low expression groups. Cut-off values used in analysis: SEMA7A in all breast cancers(all) 60mo: 113, SEMA7A all 120mo: 113, SOX9 all 60mo: 2357, SOX9 all 120mo: 1182, SEMA7A in SOX9low all 60mo: 89, SEMA7A in SOX9low all 120mo: 89, SEMA7A in SOX9high all 60mo: 118, SEMA7A in SOX9high all 120mo: 113, SEMA7A in SOX9high ER-60mo: 118, SEMA7A in SOX9high ER-120mo: 118, SEMA7A in SOX9high ER+ 60mo: 113, SEMA7A in SOX9high ER+ 120mo: 112. For gene co-expression analysis, genes were divided based on median expression. For segmentation of breast cancers based on ER status, ER status as determined by array was used.

### Statistics and Graphing

Unpaired two-tailed t-test, ordinary one-way ANOVA, and Two-way ANOVA with Sidak’s Multiple comparisons test were performed in GraphPad Prism (Dotmatics). All *in vitro* studies were at a minimum performed in biological duplicates or triplicates and technical triplicates. *In vivo* expression of markers in tissues was analyzed in at least 3 different mice or human samples, when sample availability allowed. Bar graphs represent the mean with error bars representing the mean ± SD. Box plots are median (black line), ± upper and lower quartiles (colored box), and maximum and minimum values (whiskers). Significant *p* values are represented as: \**p* ≤ 0.05, \*\**p* ≤ 0.01, \*\*\**p* ≤ 0.001, \*\*\*\**p* ≤ 0.0001. Data presented as “fold change” were normalized to controls. Graphs were generated using GraphPad Prism v10.6.1 (Dotmatics), unless otherwise specified. **Supplemental Fig. 1A and 5A** were created in BioRender Elder, A. (2025) https://BioRender.com/o5y4zgv.

## Declaration of interests

Lauren M Cozzens, Brendan Hinckley, Alan M Elder, Veronica M Wessells, Sonali Jindal, Pepper J Schedin – no conflict of interest to disclose.

Virginia F Borges – Cofounder and Chief Medical Officer of Pearl Scientific Inc., Consultant agreements with Gilead and Olema, research funding to my institution for clinical trials from Olema, Gilead, Pfizer, AstraZeneca, Abbvie, funds for advisory role from Pfizer, Gilead, AstraZeneca, Olema, and travel support from Olema.

Traci R Lyons – Cofounder and Chief Scientific Officer of Pearl Scientific Inc and Scientific Advisor, Global Cancer Therapeutics.

## AUTHOR CONTRIBUTIONS

Conception and design: L.C. and T.L.

Development of methodology: L.C, B.H., S.J., A.E., P.S., V.B., and T.L.

Acquisition of data (provided animals, acquired and managed patient samples, provided facilities, etc.): L.C, B.H., S.J., A.E., P.S., V.B., and T.L.

Analysis and interpretation of data (e.g., statistical analysis, biostatistics, computational analysis): L.C., A.E., B.H., and T.L.

Writing, review, and/or revision of the manuscript: L.C. and T.L.

Administrative, technical, or material support (i.e., reporting or organizing data, constructing databases): L.C, B.H., S.J., A.E., and T.L.

Study supervision: L.C. and T.L.

## ACKNOWLEDGEMENTS

This work was supported by NIH/NICHD R01 HD108335 (to T.R.L.), CU Anschutz Department of Medicine Aspire Program (to T.R.L. and V.F.B.), UCCC Men for the Cure (to T.R.L and V.F.B.), NIH/NCI R01 1CA169175 (V.F.B. and P.J.S.), NIH NRSA T32CA 190216-07 (to L.M.C.). *This study was supported in part by the NIH P30CA046934-funded Shared Resources:* Flow Cytometry (RRID: SCR_022035) for cytometer use and assistance, especially Alistaire Acosta for FACS assistance; Genomics (RRID: SCR_021984) for scRNA-seq; Bioinformatics (RRID: SCR_021983), especially Michael Kaufman, PhD for scRNA-seq analysis and figure generation; Human Immune Monitoring (RRID: SCR_021985), especially Troy Schedin for mIHC, and Functional Genomics (RRID: SCR_021987) for lentivirus preparation. We would also like to acknowledge the CU Anschutz Office of Laboratory Animal Research (OLAR). The authors thank the patients and their families who donated their tissue. The authors would also like to thank Weston Porter, PhD for the donation of the HC-11 cells and Steven Anderson, PhD for the donation of the EpH4 cells used in these studies. Lastly, the authors would like to thank the following for their support of this project: Heather Fairchild, PhD Rachel Steinmetz, PhD, Kelsey Kines, PhD, Taylor Rutherford, PhD, Petra Dahms, PhD, Alexandria Becks, Benjamin Bitler, PhD, Jennifer Richer, PhD, Carol Sartorius, PhD, Jenifer Monks, PhD, and Hatim Sabaawy, MD, PhD. Contents are the authors’ sole responsibility and do not necessarily represent the official NIH views.

