## Supplemental Figures for "SOX9 and SEMA7A regulate cell plasticity in the postpartum mammary gland with implications for breast cancer"

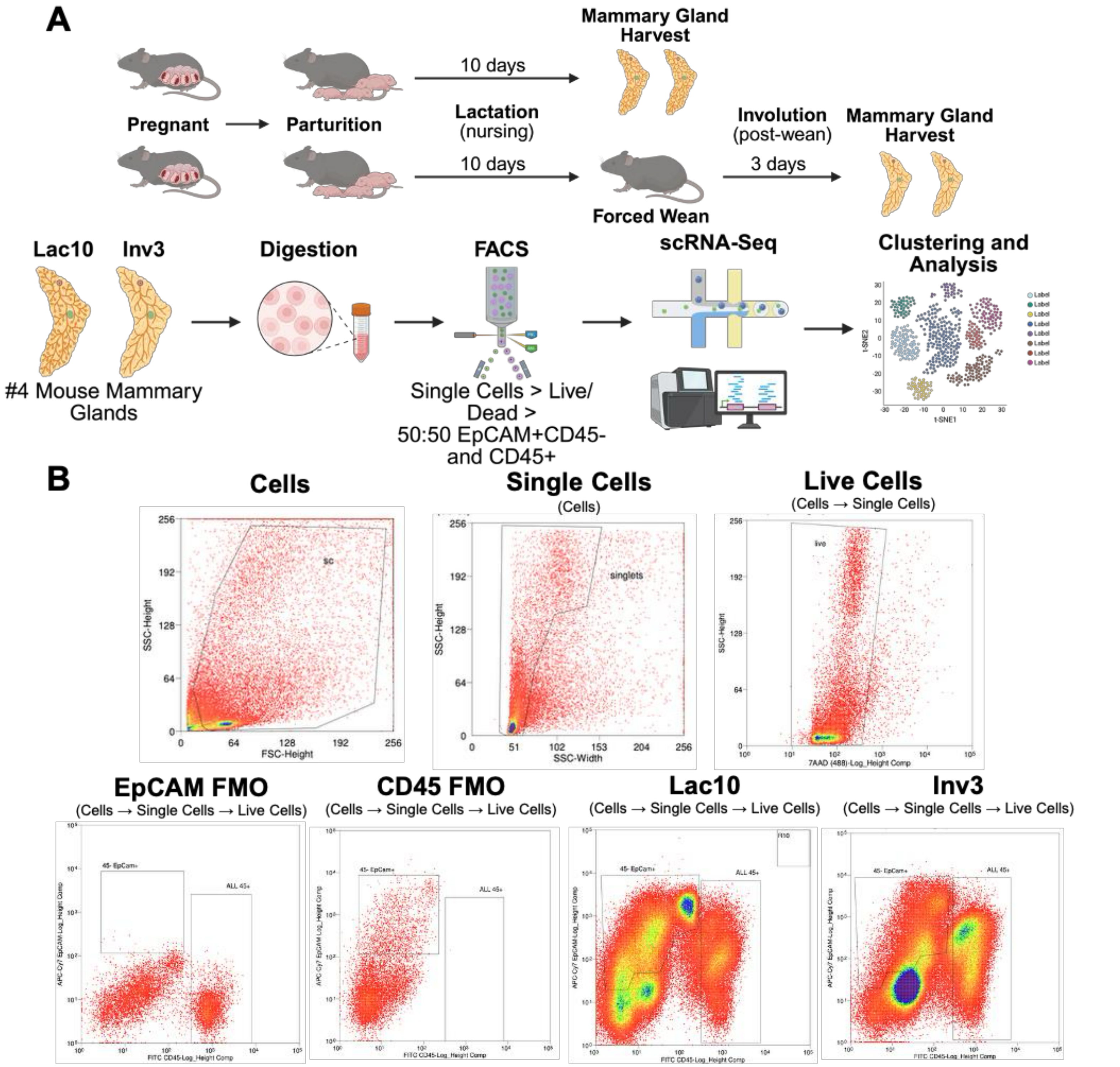


#### Supplemental Figure 1

**Supplemental Fig. 1 scRNA-seq sample collection and FACS gating strategy. A** Schematic of whole mammary gland isolation from mice undergoing lactation and involution, followed by gland digestion, FACS, capture, scRNA-seq, and analysis. **B** FACS gating strategy to isolate equal proportions of CD45+ (immune) and CD45-EpCAM+ (non-immune) live, single cells for downstream sequencing analysis.

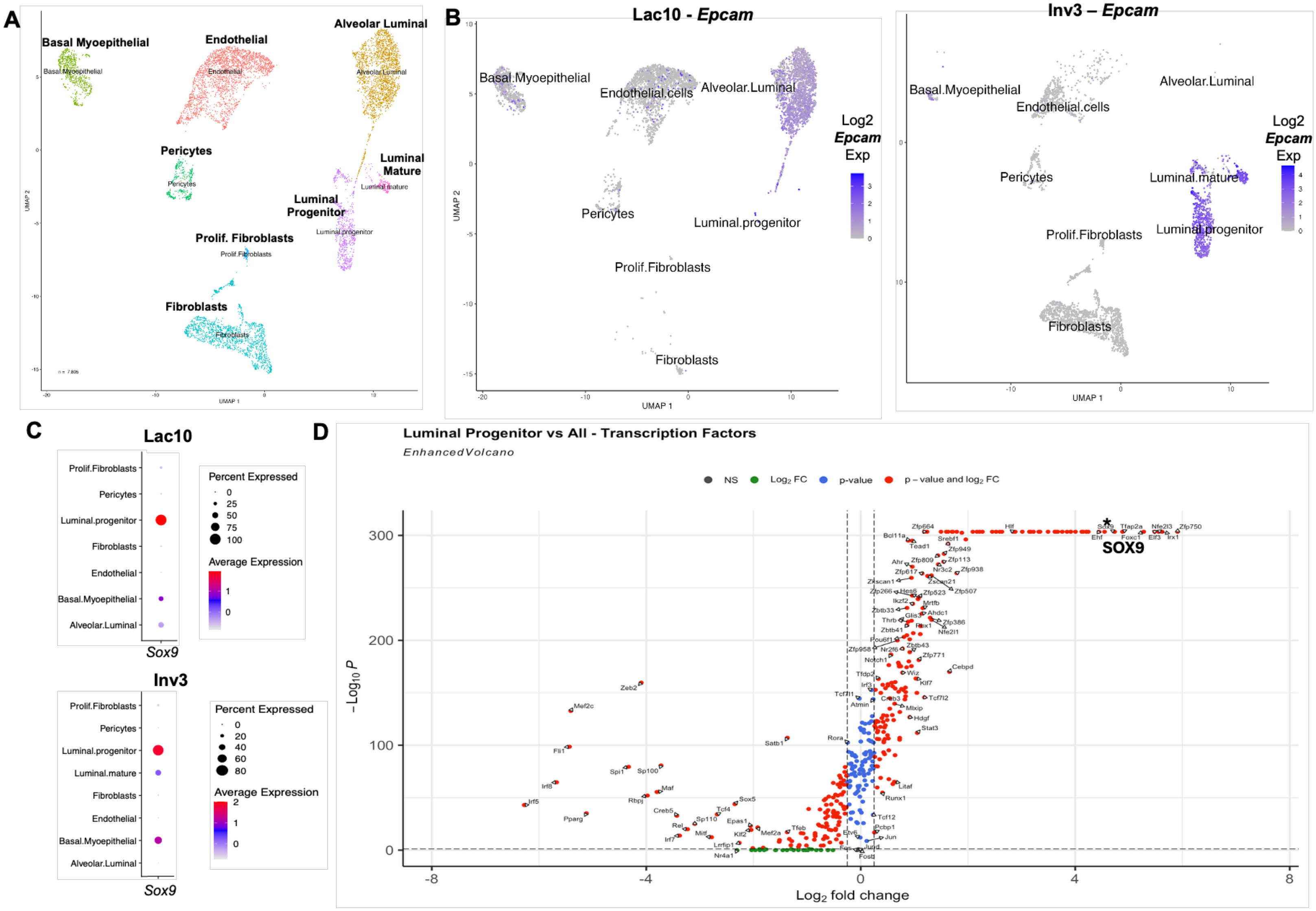


**Supplemental Figure 2**

**Supplemental Fig. 2 scRNA-seq clustering annotations and marker expression. A** UMAP visualization of 8 non-immune cell clusters from combined C56BL/6 murine mammary glands, Lac10, and Inv3 samples, with associated annotations of each cluster cell type. (n = 2 pooled #4 MGs from 1 female mouse per time point). **B** UMAP visualization of non-immune populations from Lac10 (left) and Inv3 (right) samples with overlayed Log2 *Epcam* expression. **C** Dot plot of Log2 *Sox9* expression in non-immune cell populations from Lac10 (top) or Inv3 (bottom) samples. **D** Volcano plot of transcription factor genes differentially expressed in the LPC population compared to all other cell populations from combined Lac10 and Inv4 time points. *Denotes SOX9 as a highly enriched gene in the Luminal Progenitor population.

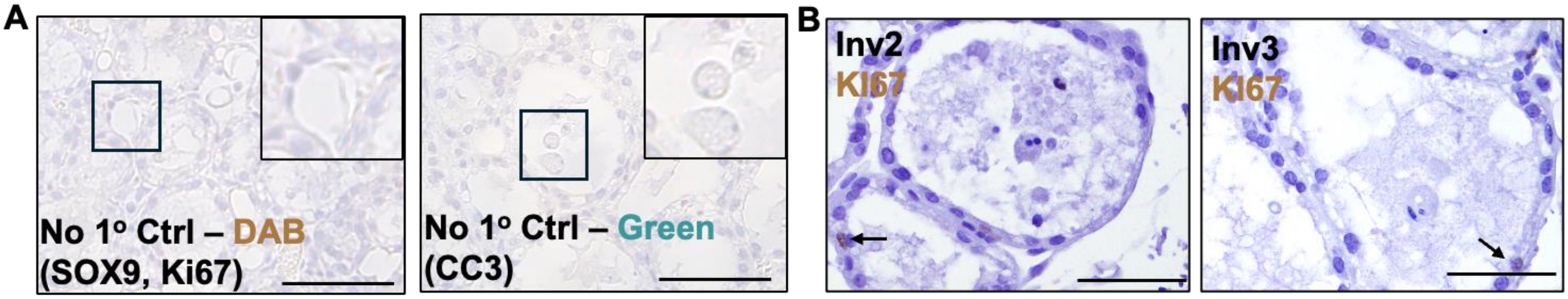


**Supplemental Figure 3**

**Supplemental Fig. 3 Murine IHC negative controls and Ki67 staining. A** Representative, no primary antibody (1º) control IHC images of murine mammary tissue for various secondary antibodies used to visualize SOX9, CC3, and Ki67. **B** Representative IHC images mouse mammary tissues stained for Ki67 Inv2 (left) and Inv3 (right) (Inv2 n = 2, Inv3 n = 2). Scale bars: 50*μ*m.

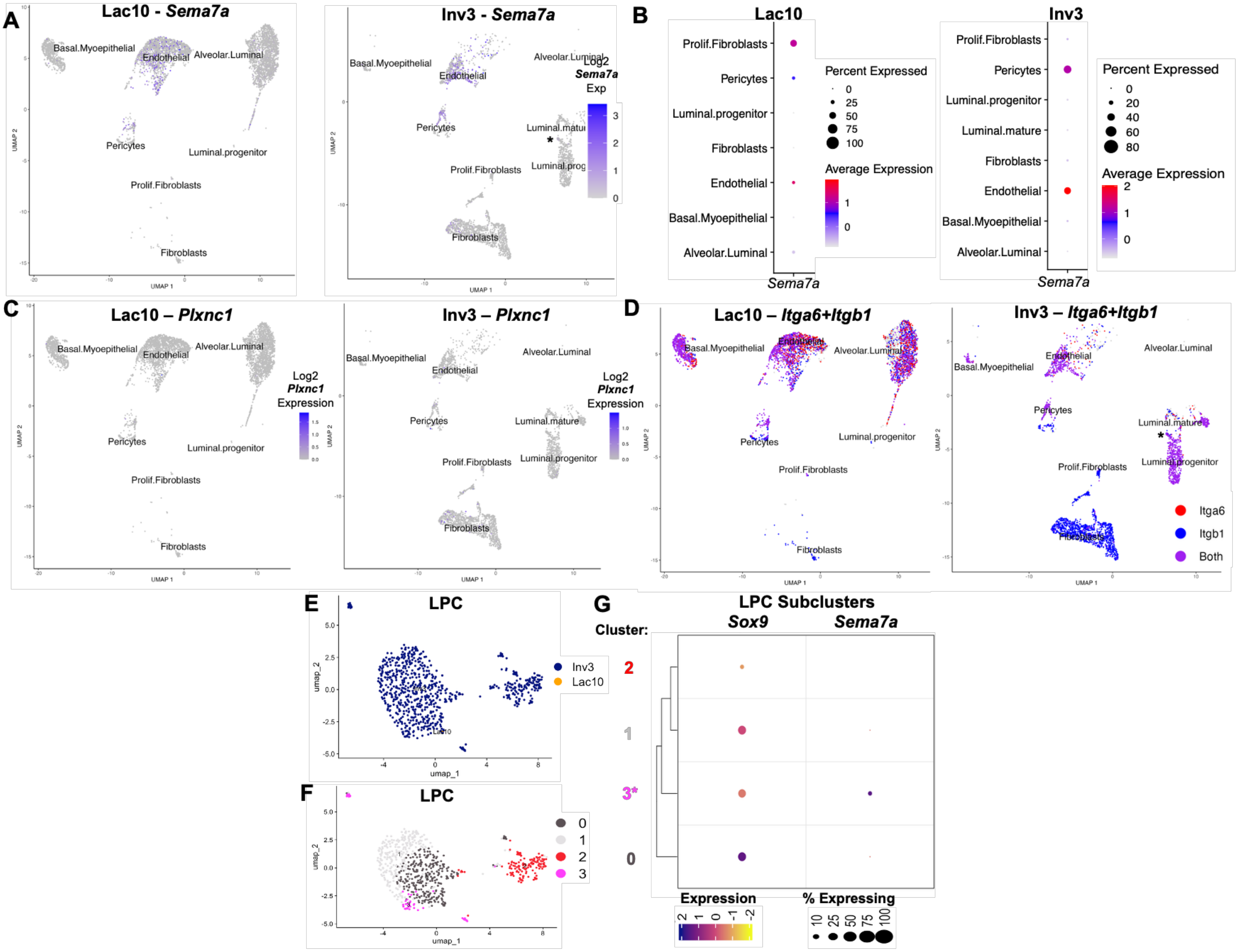


**Supplemental Figure 4**

**Supplementary Fig. 4 Integrin receptors for SEMA7A are expressed in cells of the mammary gland during lactation and involution, and sub-clustering of LPC populations. A** UMAP visualization of non-immune populations from scRNA-seq of murine mammary glands from Lac10 (left) and Inv3 (right) with overlayed Log2 *Sema7a* mRNA expression level. **B** Dot plot of Log2 *Sema7a mRNA* expression percent and level in non-immune cell populations from Lac10 (left) or Inv3 (right) samples. **C+D** UMAP visualization of Log2 expression of *Pxnc1* (**C**) and co-expression of *Itga6* and *Itgb1* (**D**) in non-immune scRNA-seq cell clusters from Lac7 (left) and Inv (right) samples. **E** Visualization of individual cell sample source (Lac10 or Inv3) for subclustered LPC populations. **F** UMAP visualization of subculturing analysis of the LPC population from combined Lac10 and Inv3 time points, showing 4 unique clusters. **G** Log2 *Sox9* or *Sema7a* expression levels in each LPC sub-cluster, * denotes the sub-cluster with both *Sox9 and Sema7a* expression.

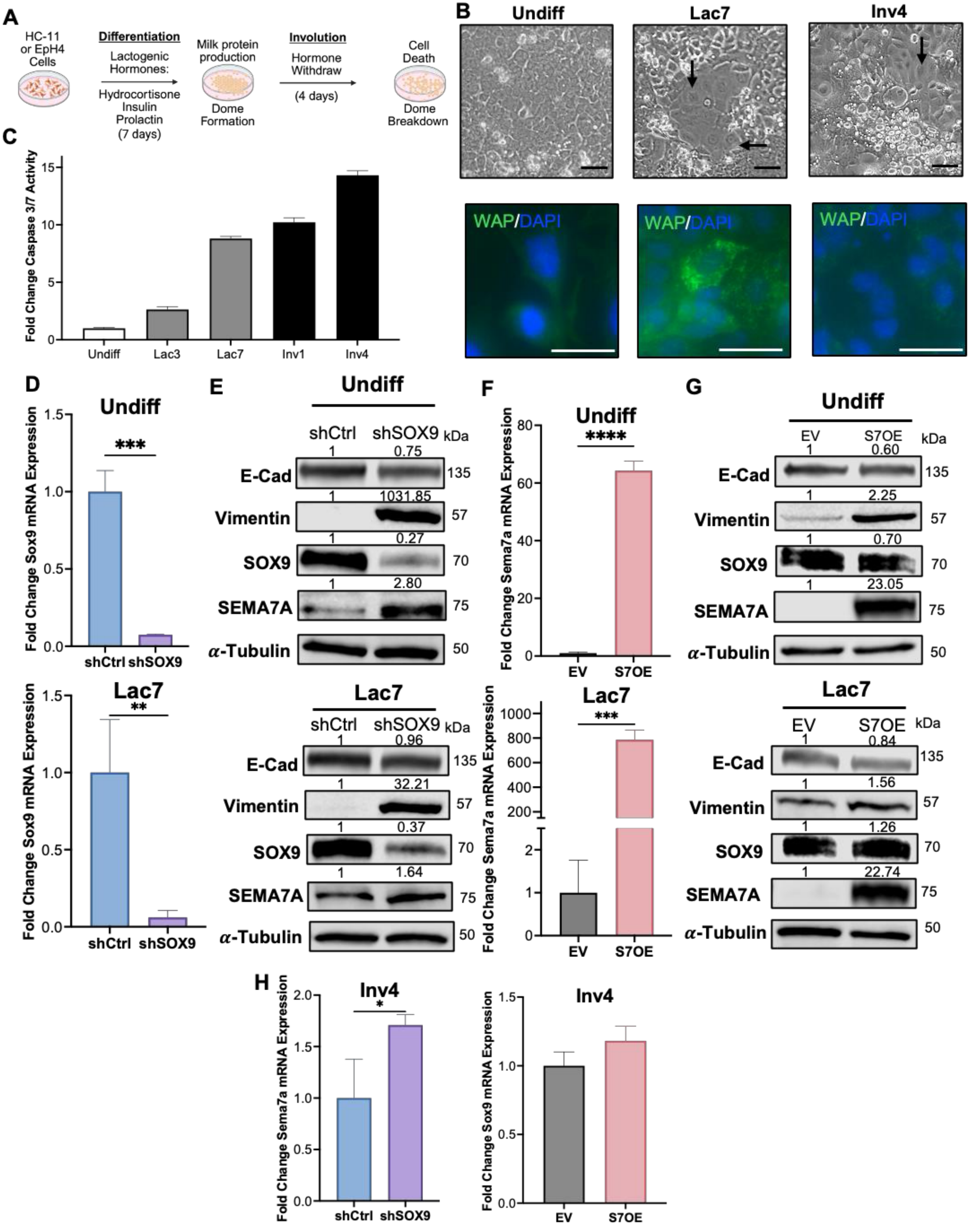


### Supplemental Figure 5

**Supplemental Fig. 5 Knockdown of *Sox9* or overexpression of *Sema7a* promotes mesenchymal cell phenotypes in undifferentiated HC-11 cultures and models of lactation. A** Schematic of *in vitro* lactogenic differentiation and involution (LacDiff/Inv) for HC-11 and EpH4 MECs. **B** Brightfield (top) image of HC-11 cell cultures across LacDiff/Inv, black arrows indicate dome formations, and immunofluorescence (bottom) of HC-11 cells for Whey Acetic Protein (WAP, green) across LacDiff/Inv; nuclei stained with DAPI (blue). Scale bars = 20$\mu$m. **C** Caspase activity assay on HC-11 cultures at indicated LacDiff/Inv time points, represented as fold change caspase 3/7 activity relative to Undiff cells (representative plots from n = 2 independent experiments with technical triplicates). **D-G** All Undiff (top) samples were collected from undifferentiated cell cultures, and all Lac7 (bottom) samples were collected on *in vitro* Lac day 7. **D+F** qPCR for *Sox9* mRNA expression in HC-11 cells harboring an shCtrl or sh*Sox9* (shSOX9) shRNA construct (**D**), or *Sema7a* mRNA expression in HC-11 harboring an EV or S7OE vector (F) represented as fold change mRNA expression relative to shCtrl (representative plot from n = 2 independent experiments with technical triplicates). **E+G** Immunoblot for SOX9, SEMA7A, E-Cadherin, and Vimentin protein levels in HC-11 shCtrl and shSOX9 cells(**E**), or EV and S7OE cells (**G**). Alpha tubulin ($\alpha$-Tubulin) was used as a loading control (representative blot from n = 3 independent experiments). **H** qPCR for *Sema7a* mRNA expression in HC-11 cells harboring an shCtrl or shSOX9 shRNA construct (left), or *Sox9* mRNA expression in HC-11 cells harboring an EV or S7OE vector (right), represented as fold change mRNA expression relative to shCtrl or EV controls, respectively. (representative plots from n = 2 independent experiments with technical triplicates). RNA was extracted on Inv day 4. Error bars represent the mean $\pm$ SD. Quantification of immunoblots represented as fold change relative to control cell cultures. Unpaired two-tailed t-test, no asterisk $p > 0.05$ *$p \leq 0.05$, **$p \leq 0.01$, ***$p \leq 0.001$, ****$p \leq 0.0001$.

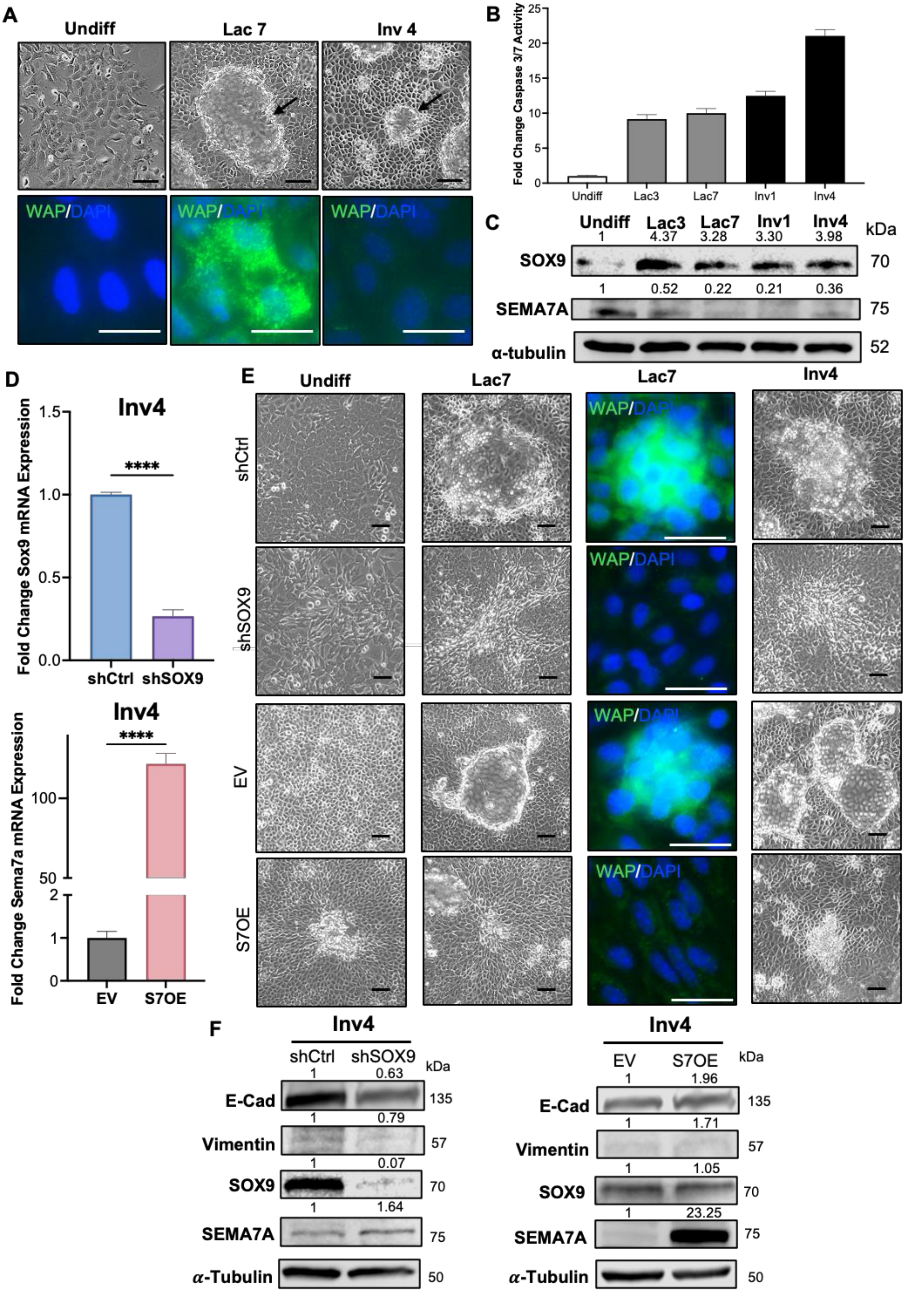


### Supplemental Figure 6

**Supplemental Fig. 6 Knockdown of *Sox9* or overexpression of *Sema7a* impairs differentiation and promotes mesenchymal cell phenotypes in EpH4 *in vitro* models of lactation and involution. A** Brightfield images (top) of EpH4 cell cultures across LacDiff/Inv, black arrows indicate dome formations, and immunofluorescence images (bottom) of EpH4 cells for Whey Acetic Protein (WAP, green) across LacDiff/Inv; nuclei stained with DAPI (blue). Scale bars = 20$\mu$m. **B** Caspase activity assay on EpH4 cultures at indicated LacDiff/Inv time points, represented as fold change caspase 3/7 activity relative to Undiff cells (representative plots from n = 2 independent experiments with technical triplicates). **C** Immunoblot showing protein levels of SOX9 and SEMA7A in undifferentiated (Undiff), early (Lac3), and late lactogenic differentiation (Lac7), and early (Inv1) and late involution (Inv4) in Wildtype (WT) EpH4 cell cultures (representative blot of n = 3 independent experiments). Alpha tubulin ($\alpha$-Tubulin) was used as a loading control. **D** qPCR for *Sox9* mRNA expression in EpH4 cells harboring an shCtrl or shSOX9 shRNA construct (top), or *Sema7a* mRNA expression in EpH4 cells harboring an EV or S7OE vector (bottom), represented as fold change mRNA expression relative to shCtrl or EV controls, respectively. (representative plots from n = 2 independent experiments with technical triplicates). RNA was extracted on Inv day 4. **E** Brightfield images of EpH4 shCtrl+shSOX9 (top) and EV+S7OE (bottom) cultures across LacDiff/Inv, black arrows indicate dome formation (representative blot from n = 3 independent experiments). Immunofluorescence of EpH4 shCtrl+shSOX9 (top) and EV+S7OE (bottom) cells for Whey Acetic Protein (WAP, green) on Lac7; nuclei stained with DAPI (blue). Scale bars = 20$\mu$m. **F** Immunoblot for SOX9, SEMA7A, E-Cadherin, and Vimentin protein levels in EpH4 shCtrl+shSOX9 (left) and EV+S7OE (right) cells. $\alpha$-Tubulin was used as a loading control (representative blot from n = 3 independent experiments). All samples collected on Inv day 4. Error bars represent the mean $\pm$ SD. Quantification of immunoblots represented as fold change relative to control or undifferentiated cell cultures. Unpaired two-tailed t-test, ****$p \leq 0.0001$.

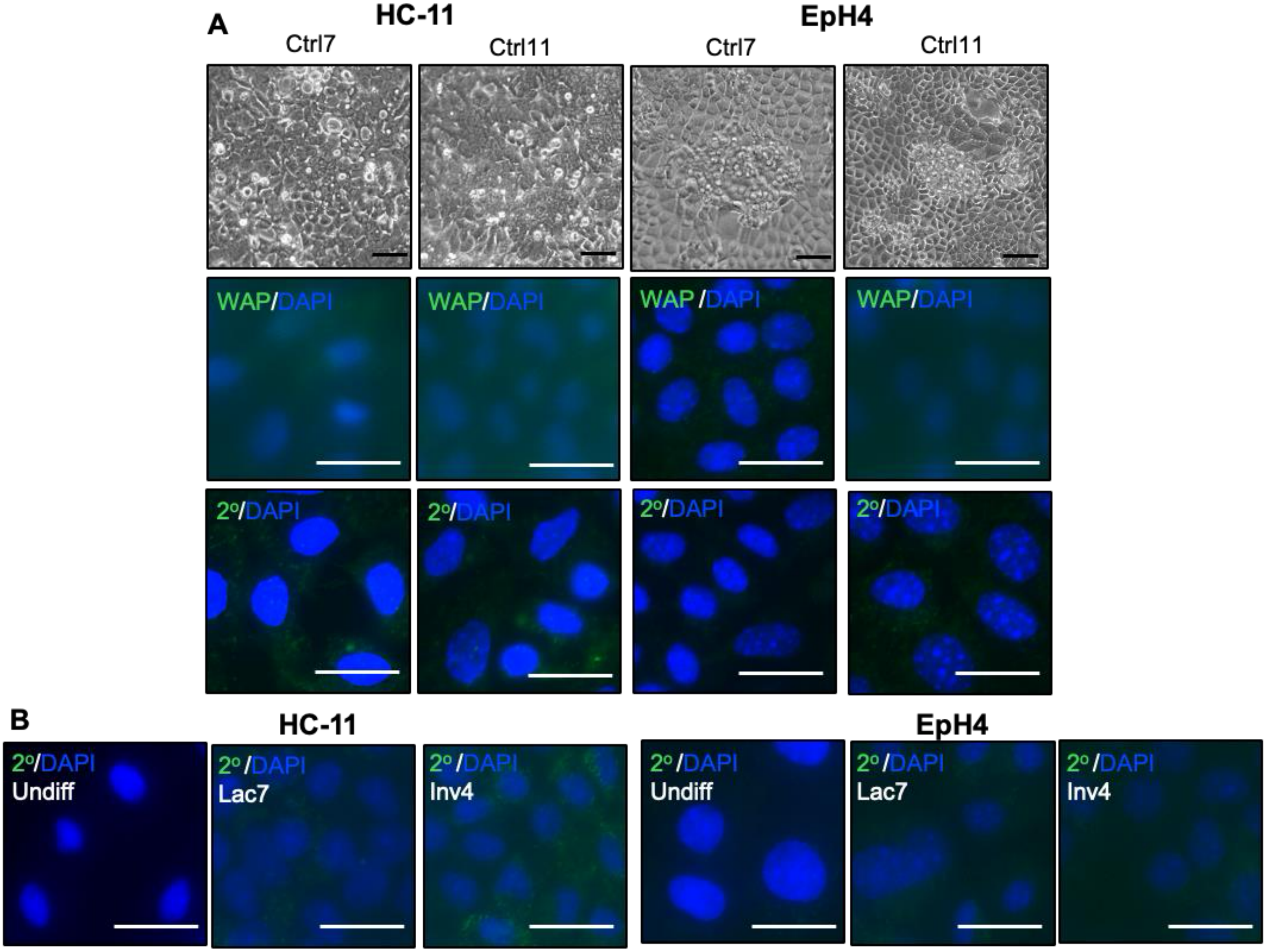


#### Supplemental Figure 7

**Supplemental Fig. 7 Dome formation and milk protein production in murine MEC cultures are dependent on treatment with lactogenic hormones. A** Top: brightfield images of HC-11 (left) and EpH4 (right) cell cultures maintained in control prime media without lactogenic hormones for 7 and 11 days (Ctrl7+11 corresponding to Lac7 and Inv4 in Differentiation cultures). Middle: IF in HC-11 and EpH4 Ctrl cells for Whey Acetic Protein (WAP, green; DAPI, blue), Bottom: no primary (1º) antibody secondary only controls for Ctrl cells, secondary antibody only (2º antibody, green; DAPI, blue). **B** No 1º antibody secondary only controls for HC-11 (left) and EpH4 (right) Undiff and LacDiff/Inv cells (2º antibody, green; DAPI, blue). **A+B** Scale bars = 20$\mu$m.

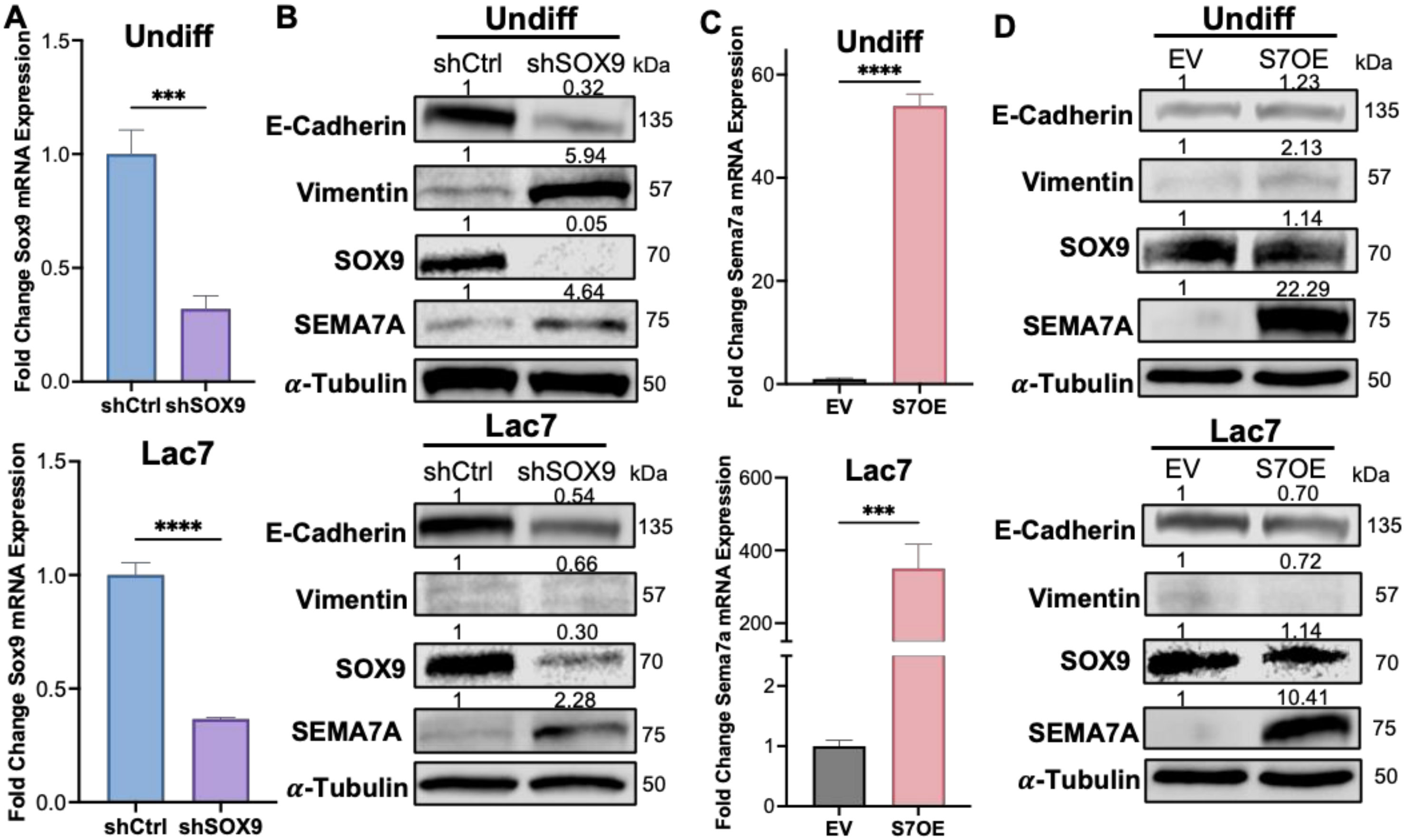


#### Supplemental Figure 8

**Supplemental Fig. 8 Knockdown of *Sox9* or overexpression of *Sema7a* promotes mesenchymal cell phenotypes in undifferentiated EpH4 cultures and models of lactation. A-D** All Undiff (top) samples were collected from undifferentiated cell cultures, and all Lac7 (bottom) samples were collected on *in vitro* Lac Day 7. **A** qPCR for *Sox9* mRNA expression in EpH4 cells harboring an shCtrl or shSOX9 construct, represented as fold change mRNA expression relative to shCtrl (representative plot from n = 2 independent experiments with technical triplicates). **B** Immunoblot for SOX9, SEMA7A, E-Cadherin, and Vimentin protein levels in EpH4 shCtrl and shSOX9 cells. Alpha tubulin was used as a loading control (representative blot from n = 3 independent experiments). **C** qPCR for *Sema7a* mRNA expression in EpH4 harboring an EV or S7OE vector, represented as fold change mRNA expression relative to EV (representative plot from n = 2 independent experiments with technical triplicates). **D** Immunoblot for SOX9, SEMA7A, E-Cadherin, and Vimentin protein levels in EpH4 EV and S7OE cells. Alpha tubulin was used as a loading control (representative blot from n = 3 independent experiments). Quantification of immunoblots represented as fold change relative to control cell cultures Error bars represent the mean ± SD. Unpaired two-tailed t-test, *** $p \leq 0.001$, ****$p \leq 0.0001$.

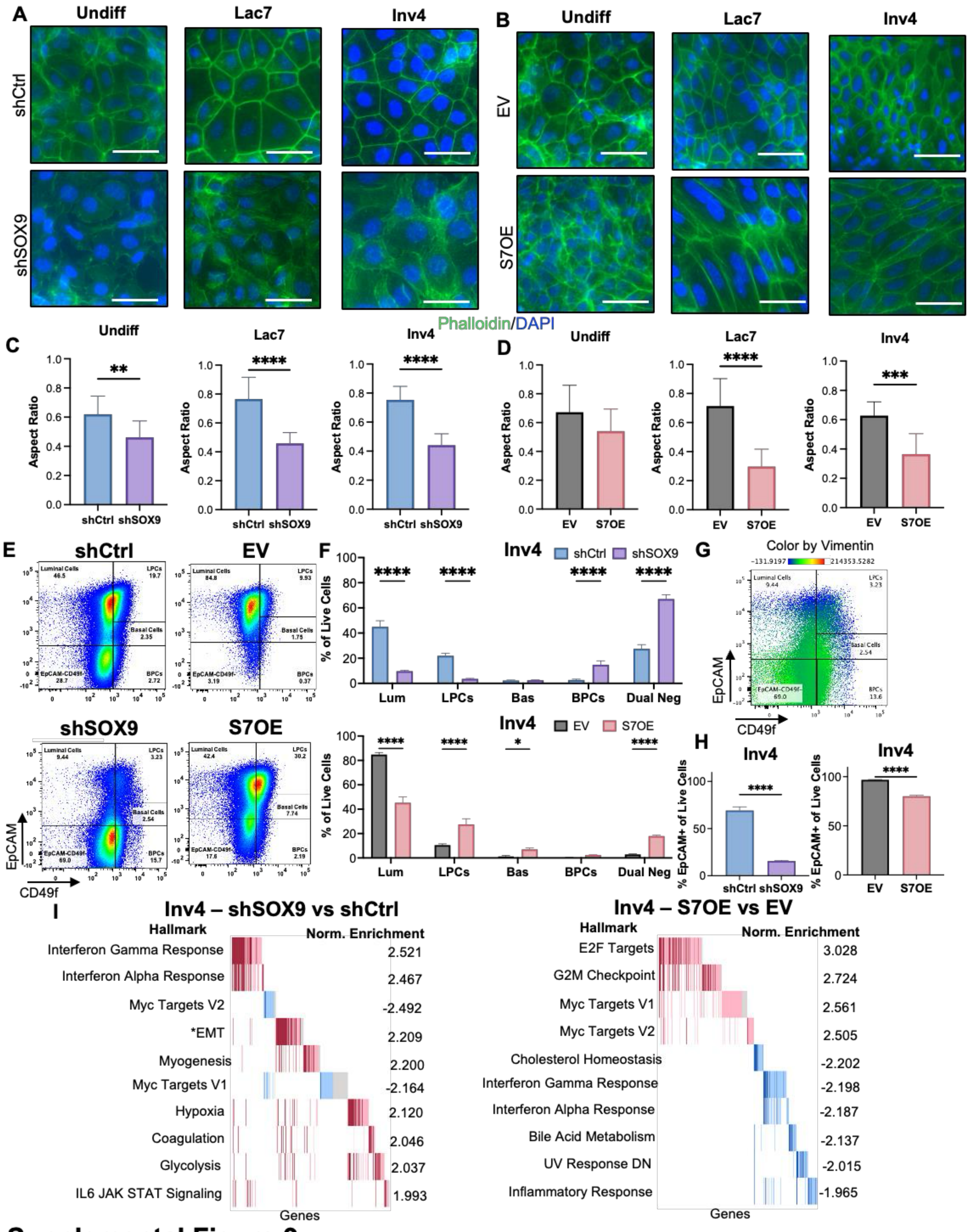


### Supplemental Figure 9

**Supplemental Fig. 9 Loss of SOX9 or gain of SEMA7A promotes the emergence of a mesenchymal cell state in EpH4 models of involution. A+B** Phalloidin stain images of EpH4 shCtrl+shSOX9 (A) and EV+S7OE (B) cultures across LacDiff/Inv (representative photos from n = 2 independent experiments). Scale bars = 20$\mu$m. **C+D** Aspect ratio calculations (width/length) from phalloidin staining in A+B (n = 10 cells/group). **E** Representative flow cytometry plots of EpCAM (y-axis) and CD49f (x-axis) staining in EpH4 cultures to visualize MEC subtypes, shCtrl+shSOX9 (left) and EV+S7OE (right). Numbers denote the percentage of that subtype of all live cells (representative plots of n = 3 independent experiments with technical triplicates). **F** Quantification of percent individual MEC subtypes from flow cytometry of all live cells in EpH4 shCtrl+shSOX9 cells (top) and EV+S7OE cells (bottom). **G** Example flow cytometry plot of EpCAM (y-axis) and CD49f (x-axis) staining, colored by Vimentin fluorescence intensity. **H** Quantification of percent EpCAM+ cells from flow cytometry of all live cells in EpH4 shCtrl+shSOX9 cells (left) and EV+S7OE cells (right). **I** Top 10 most significantly (adj. p-value) enriched Hallmark pathways from RNA-seq of EpH4 shSOX9 (left) or S7OE (right) Inv4 cultures compared to their associated controls, shCtrl and EV, respectively. Normalized enrichment values shown. *denotes epithelial to mesenchymal transition (EMT) as a top pathway for shSOX9 cultures (n = 3 biological triplicates). Error bars represent the mean $\pm$ SD. **C+H** Unpaired two-tailed t-test. **F** Two-way ANOVA Sidak's Multiple comparisons test. No asterisk $p > 0.05$, *$p \leq 0.05$, **$p \leq 0.01$, ***$p \leq 0.01$ ****$p \leq 0.0001$.

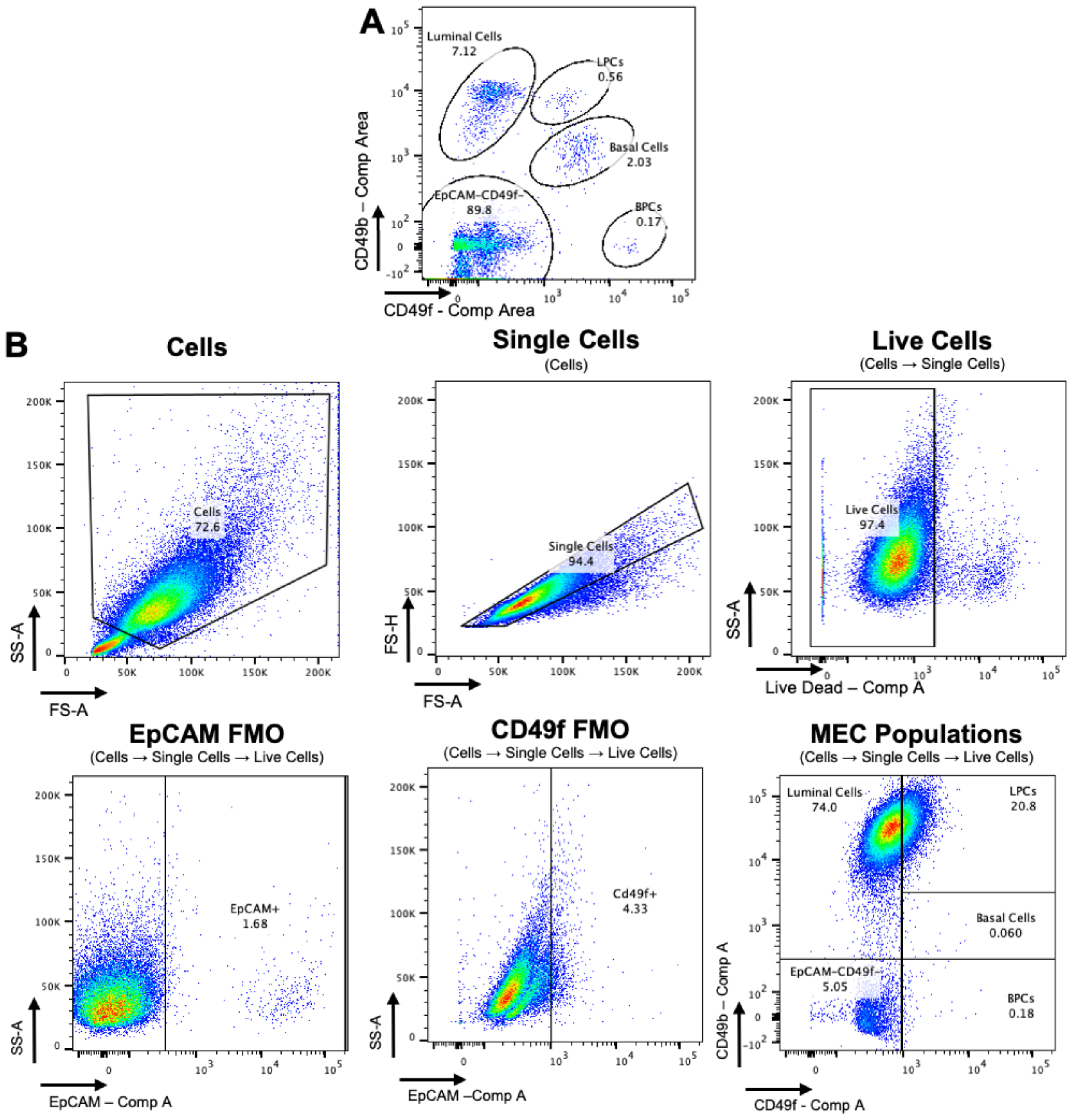


**Supplemental Figure 10**

**Supplemental Fig. 10 MEC Subtype Flow Cytometry Gating Strategy for *in vitro* cultures. A** Example of MEC populations isolated from murine mammary glands for which this EpCAM by CD49f gating strategy was originally developed. **B** For all flow cytometry experiments, cells were identified by plotting side scatter area by forward scatter area (SS-A and FS-A) to remove debris. Single cells were then identified by plotting FS height (FS-H) by FS-A. Live cells were identified by Live/Dead viability staining. FMO controls were used on live cells to determine EpCAM and CD49f thresholds for MEC subtype gating. Gates were validated with single-stain controls and unstained cell controls of the same cell type. Luminal cells (Lum - $EpCAM^{+}CD49f^{-}$), luminal progenitor cells (LPC - $EpCAM^{+}CD49f^{+}$), basal cells (Bas - $EpCAM^{low/mid}CD49f^{+}$), basal progenitor cells (BPC - $EpCAM^{-}CD49f^{+}$), dual negative cells (Dual Neg – $EpCAM^{-}CD49f^{-}$). Comp A = compensated fluorescence signal area.

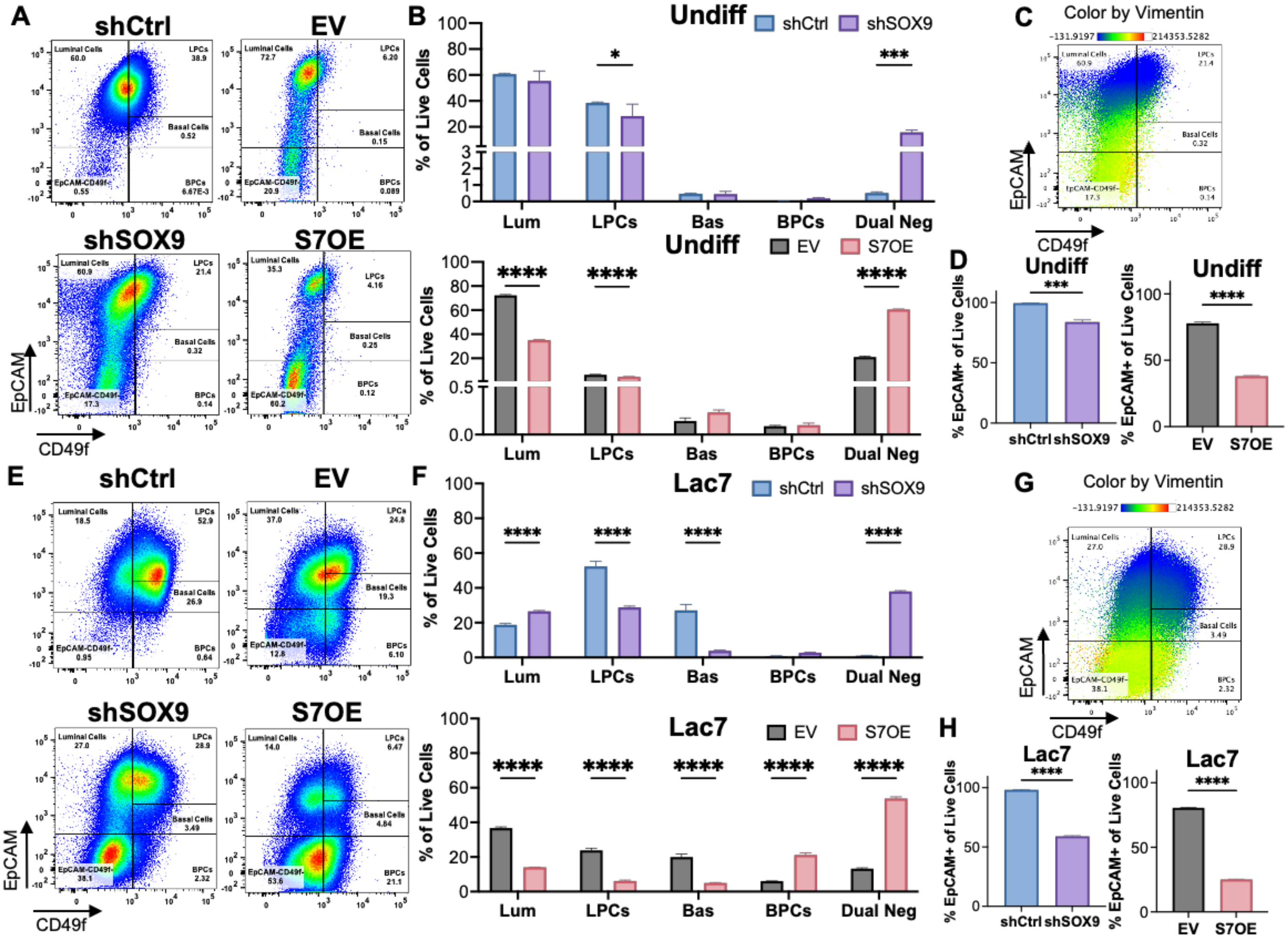


**Supplemental Figure 11**

**Supplemental Fig. 11 Loss of SOX9 or gain of SEMA7A promotes the emergence of a mesenchymal cell state in undifferentiated HC-11 cultures and models of lactation. A+E** Representative flow cytometry plots of EpCAM (y-axis) and CD49f (x-axis) staining in undifferentiated HC-11 cultures (**A**) and Lac7 HC-11 cultures (**E**) to visualize MEC subtypes, shCtrl+shSOX9 (left) and EV+S7OE (right). Numbers denote the percentage of that subtype of all live cells (representative plots of n = 3 independent experiments with technical triplicates). **B+F** Quantification of percent individual MEC subtypes from flow cytometry of all live cells in undifferentiated HC-11 cultures (**B**) and Lac7 HC-11 cultures (**F**) shCtrl+shSOX9 cells (top) and EV+S7OE cells (bottom). **C+G** Example flow cytometry plot of EpCAM (y-axis) and CD49f (x-axis) staining, colored by Vimentin fluorescence intensity in undifferentiated HC-11 cultures (**C**) and Lac7 HC-11 cultures (**G**). **D+H** Quantification of percent EpCAM+ cells from flow cytometry of all live cells in undifferentiated HC-11 cultures (**D**) and Lac7 HC-11 cultures (**H**) shCtrl+shSOX9 cells (left) and EV+S7OE cells (right). Error bars represent the mean ± SD. **B+F** Two-way ANOVA Sidak's Multiple comparisons test. **D+H** Unpaired two-tailed t-test. No asterisk p > 0.05, *$p \leq 0.05$, ****$p \leq 0.0001$.

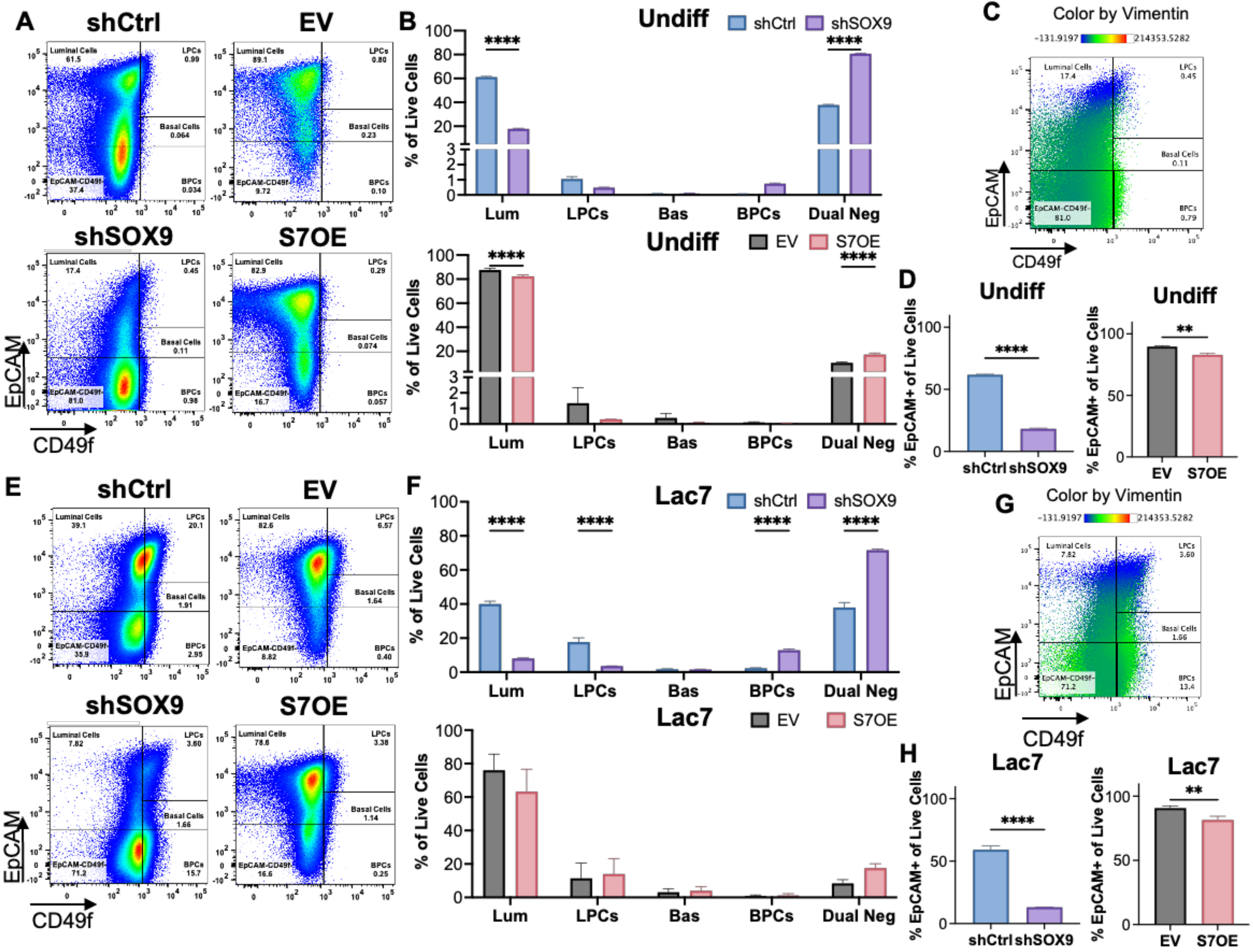


**Supplemental Figure 12**

**Supplemental Fig. 12 Loss of SOX9 or gain of SEMA7A promotes the emergence of a mesenchymal cell state in undifferentiated EpH4 cultures and models of lactation. A+E** Representative flow cytometry plots of EpCAM (y-axis) and CD49f (x-axis) staining in undifferentiated EpH4 cultures (**A**) and Lac7 EpH4 cultures (**E**) to visualize MEC subtypes, shCtrl+shSOX9 (left) and EV+S7OE (right). Numbers denote the percentage of that subtype of all live cells (representative plots of n = 3 independent experiments with technical triplicates). **B+F** Quantification of percent individual MEC subtypes from flow cytometry of all live cells in undifferentiated EpH4 cultures (**B**) and Lac7 EpH4 cultures (**F**) shCtrl+shSOX9 cells (top) and EV+S7OE cells (bottom). **C+G** Example flow cytometry plot of EpCAM (y-axis) and CD49f (x-axis) staining, colored by Vimentin fluorescence intensity in undifferentiated EpH4 cultures (**C**) and Lac7 EpH4 cultures (**G**). **D+H** Quantification of percent EpCAM+ cells from flow cytometry of all live cells in undifferentiated EpH4 cultures (**D**) and Lac7 EpH4 cultures (**H**) shCtrl+shSOX9 cells (left) and EV+S7OE cells (right). Error bars represent the mean $\pm$ SD. **B+F** Two-way ANOVA Sidak's Multiple comparisons test. **D+H** Unpaired two-tailed t-test. No asterisk $p > 0.05$, $^{**}p \leq 0.01$, $^{****}p \leq 0.0001$.

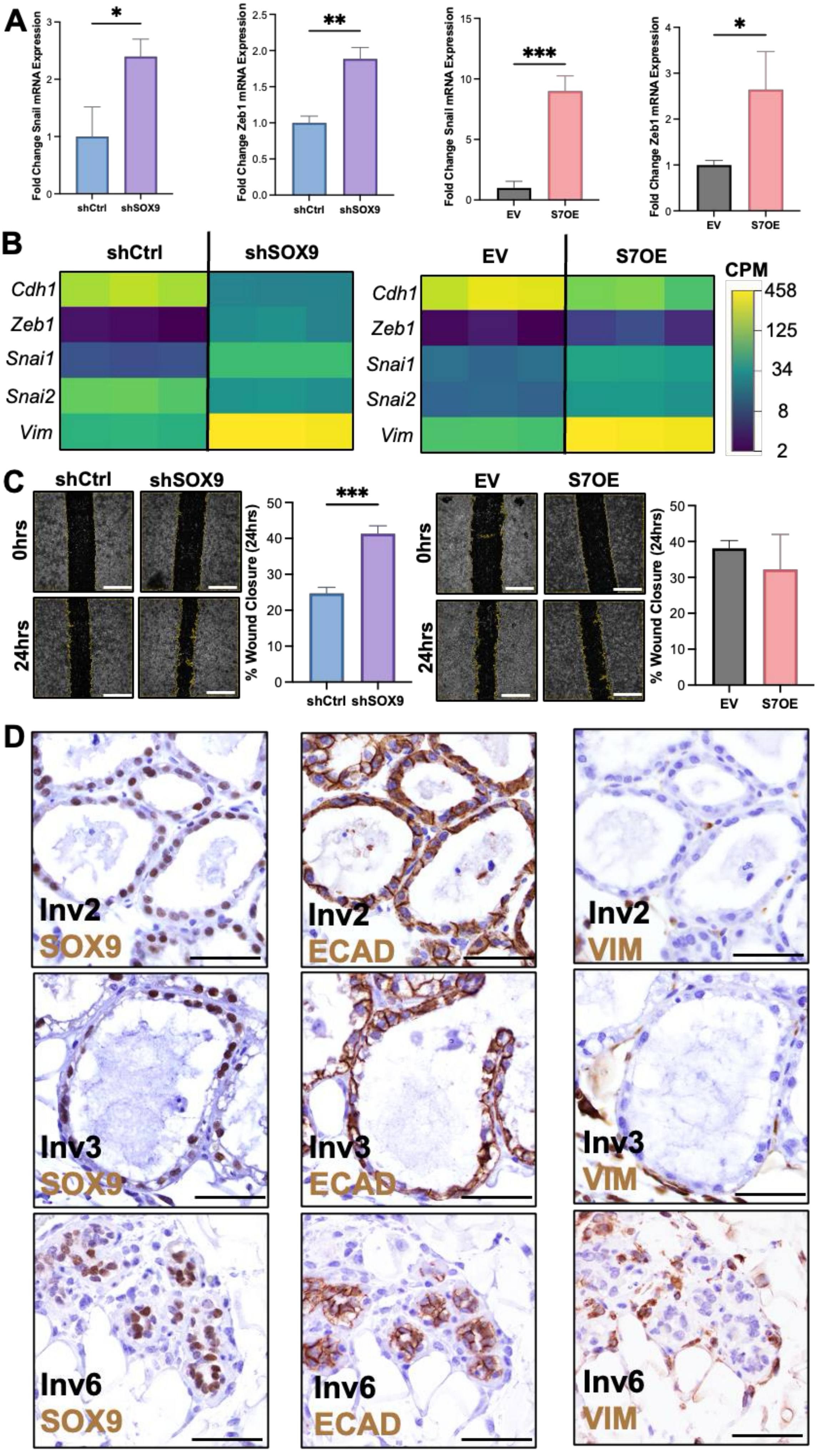


Supplemental Figure 13

**Supplemental Fig. 13 Loss of SOX9 or gain of SEMA7A promotes mesenchymal cell phenotypes in models of involution, and SOX9 is expressed in E-Cadherin+ cells during involution *in vivo*. A-C** All experiments were performed on or after *in vitro* Inv day 4. **A** qPCR for the EMT transcription factors, *Snail* or *Zeb1*, mRNA expression in shCtrl or shSOX9 (left) or EV or S7OE (right) HC-11 cells, represented as fold change mRNA expression relative to shCtrl or EV (representative plot from n = 2 independent experiments with technical triplicates). **B** mRNA expression of various EMT markers from bulk RNA-seq of HC-11 shCtrl and shSOX9 Inv4 cells in read counts per million (CPM). E-Cadherin (*Cdh1*), Zeb1, Snail (*Snai1*), Slug (*Snai2*), and Vimentin (*Vim*). **C** Representative images of HC-11 shCtrl+shSOX9 (left) and EV+S7OE (right) scratch wound closure after 24 hours. Quantification of percent wound closure after 24 hours. **C** Representative images, at a lower magnification than those presented in Fig. 4J, of serial sections of murine mammary glands from Inv day 2, 3, or 6 stained for SOX9, E-Cadherin, or Vimentin. Error bars represent the mean $\pm$ SD. Unpaired two-tailed t-test. No asterisk $p > 0.05$, *$p \leq 0.05$, **$p \leq 0.01$, ***$p \leq 0.001$.

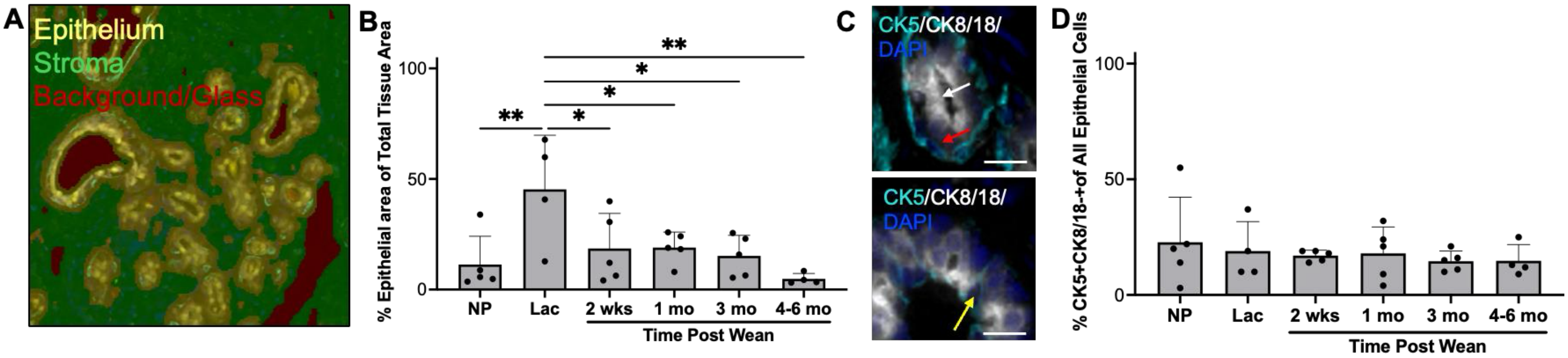


**Supplemental Figure 14**

**Supplemental Fig. 14 Epithelial tissue increases during lactation, while luminal/basal hybrid cell proportions remain consistent across reproductive stages of the human mammary gland. A** Example of Akoya Biosciences InForm software segmentation of epithelial and stromal compartments in a representative human mammary tissue sample. Epithelium is overlayed in yellow, stroma is overlaid in green, and background/glass areas where there is no tissue are overlaid in red. **B** Quantification of the percent epithelial area of total tissue area per tissue section. **C** Representative image of a CK5+CK8/18- basal cell (cyan, red arrow), a CK5-CK8/18+ luminal cell (white, white arrow), and a CK5+CK8/18+ basal/luminal hybrid cell (cyan+white, yellow arrow); nuclei stained with DAPI (blue). Scale bars: 15μm. **D** Quantification of percent basal/luminal CK5+CK8/18+ hybrid cells of all epithelial cells per tissue section. Error bars represent the mean ± SD. Ordinary one-way ANOVA Sidak's Multiple comparisons test. No asterisk $p > 0.05$, *$p \leq 0.05$, **$p \leq 0.01$.

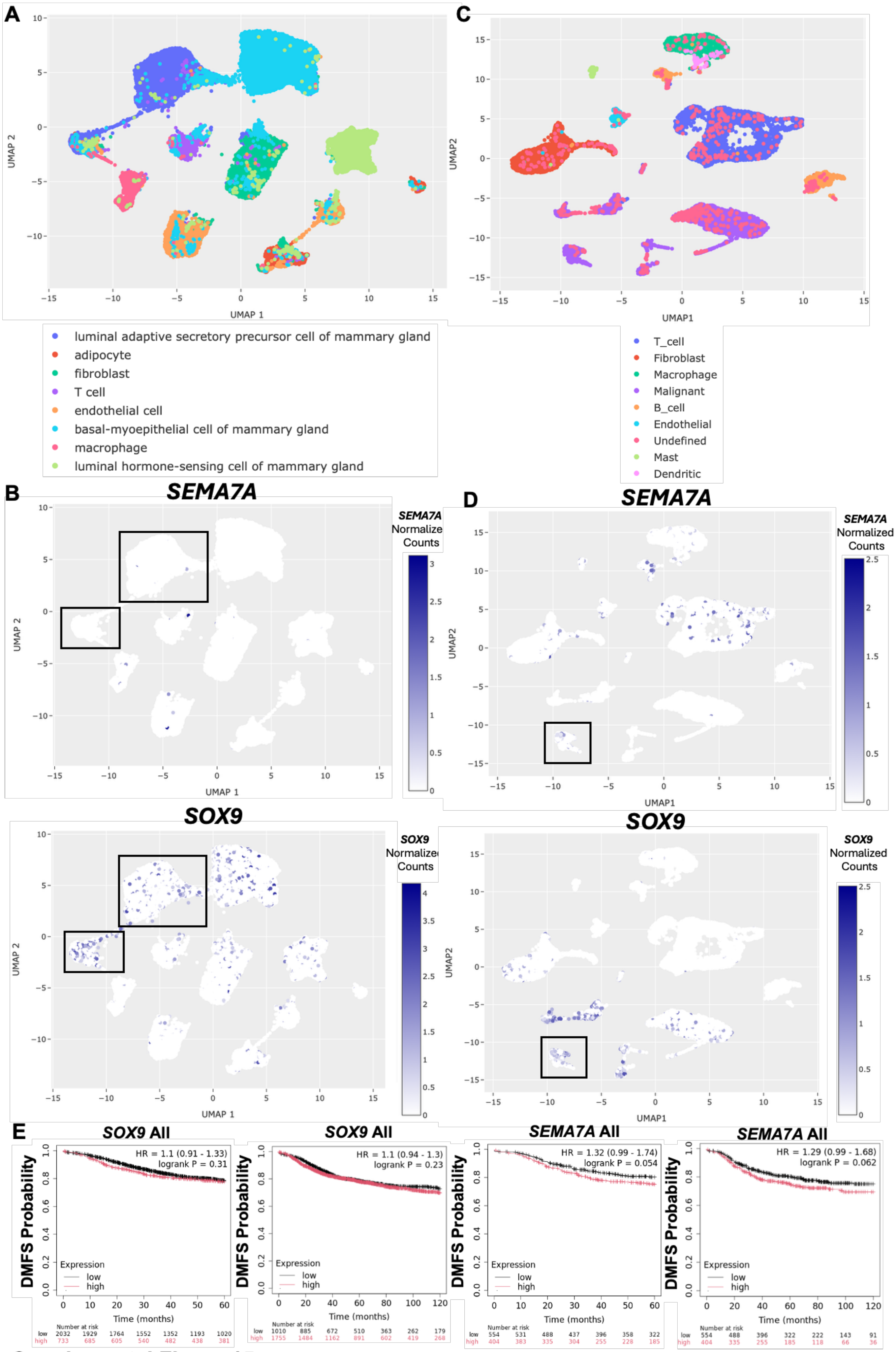


Supplemental Figure 15

**Supplemental Fig. 15 *SOX9* and *SEMA7A* are expressed in human breast epithelium and in breast tumors. A** Annotated UMAP from scRNA-seq analysis of normal human breast tissue from Bhat-Nakshatri et. al, 2024, identifying 8 unique clusters. n = 66 samples. **B** Full UMAPs overlayed with normalized counts of *SEMA7A* (top) or *SOX9* (bottom) mRNA expression. Boxes denote populations further explored in Fig. 6D. **C** Annotated UMAP from scRNA-seq analysis of human breast tumors from Qian et al., 2020, identifying 9 unique populations. n = 7 samples. **D** Full UMAPs overlayed with normalized counts of *SEMA7A* (top) or *SOX9* (bottom) mRNA expression, from Qian et. al, 2020 data set. **E** Breast cancer patient Kaplan-Meier 5-year (60-month, top) and 10-year (120-month, bottom) DMFS analysis of low and high *SOX9* (left), and low and high *SEMA7A* (right) mRNA expression in tumor samples from all subtypes of breast cancer combined from KM-plotter. Boxes denote populations further explored in Fig. 6E.
