## Supplemental Table 1 for "SOX9 and SEMA7A regulate cell plasticity in the postpartum mammary gland with implications for breast cancer"

**Supplemental Table 1: Selected cell segmentation features for Neural Network training.** Cellular segmentation features used to train the neural network to separate stromal and epithelial cells. DAB staining measurements were excluded to prevent classifying decisions based on the presence or absence of our SOX9 stain.

| **Feature** | **Selected (True/False)** |
|---|---|
| Area µm^2 | True |
| Length µm | True |
| Circularity | True |
| Solidarity | True |
| Max diameter µm | True |
| Min diameter µm | True |
| Hematoxylin: Mean | True |
| Hematoxylin: Median | True |
| Hematoxylin: Min | True |
| Hematoxylin: Max | True |
| Hematoxylin: Std.Dev | True |
| DAB: Mean | False |
| DAB: Median | False |
| DAB: Min | False |
| DAB: Max | False |
| Smoothed 25 µm: Area µm^2 | True |
| Smoothed 25 µm: Length µm | True |
| Smoothed 25 µm: Circularity | True |
| Smoothed 25 µm: Solidarity | True |
| Smoothed 25 µm: Max diameter µm | True |
| Smoothed 25 µm: Min diameter µm | True |
| Smoothed 25 µm: Hematoxylin: Mean | True |

| | |
|---|---|
| Smoothed 25 µm: Hematoxylin: Median | True |
| Smoothed 25 µm: Hematoxylin: Min | True |
| Smoothed 25 µm: Hematoxylin: Max | True |
| Smoothed 25 µm: Hematoxylin: Std.Dev | True |
| Smoothed 25 µm: DAB: Mean | False |
| Smoothed 25 µm: DAB: Median | False |
| Smoothed 25 µm: DAB: Min | False |
| Smoothed 25 µm: DAB: Max | False |
| Smoothed 25 µm: Nearby detection counts | True |
| Smoothed 50 µm: Area µm^2 | True |
| Smoothed 50 µm: Length µm | True |
| Smoothed 50 µm: Circularity | True |
| Smoothed 50 µm: Solidarity | True |
| Smoothed 50 µm: Max diameter µm | True |
| Smoothed 50 µm: Min diameter µm | True |
| Smoothed 50 µm: Hematoxylin: Mean | True |
| Smoothed 50 µm: Hematoxylin: Median | True |
| Smoothed 50 µm: Hematoxylin: Min | True |
| Smoothed 50 µm: Hematoxylin: Max | True |
| Smoothed 50 µm: Hematoxylin: Std.Dev | True |
| Smoothed 50 µm: DAB: Mean | False |
| Smoothed 50 µm: DAB: Median | False |
| Smoothed 50 µm: DAB: Min | False |
| Smoothed 50 µm: DAB: Max | False |
| Smoothed 50 µm: Nearby detection counts | True |
